# State-transition Modeling of Blood Transcriptome Predicts Disease Evolution and Treatment Response in Chronic Myeloid Leukemia

**DOI:** 10.1101/2023.10.11.561908

**Authors:** David E. Frankhouser, Russell C. Rockne, Lisa Uechi, Dandan Zhao, Sergio Branciamore, Denis O’Meally, Jihyun Irizarry, Lucy Ghoda, Haris Ali, Jeffery M. Trent, Stephen Forman, Yu-Hsuan Fu, Ya-Huei Kuo, Bin Zhang, Guido Marcucci

## Abstract

Chronic myeloid leukemia (CML) is initiated and maintained by BCR::ABL which is clinically targeted using tyrosine kinase inhibitors (TKIs). TKIs can induce long-term remission but are also not curative. Thus, CML is an ideal system to test our hypothesis that transcriptome-based state-transition models accurately predict cancer evolution and treatment response. We collected time-sequential blood samples from tetracycline-off (Tet-Off) BCR::ABL-inducible transgenic mice and wild-type controls. From the transcriptome, we constructed a CML statespace and a three-well leukemogenic potential landscape. The potential’s stable critical points defined observable disease states. Early states were characterized by anti-CML genes opposing leukemia; late states were characterized by pro-CML genes. Genes with expression patterns shaped similarly to the potential landscape were identified as drivers of disease transition. Re-introduction of tetracycline to silence the BCR::ABL gene returned diseased mice transcriptomes to a near healthy state, without reaching it, suggesting parts of the transition are irreversible. TKI only reverted the transcriptome to an intermediate disease state, without approaching a state of health; disease relapse occurred soon after treatment. Using only the earliest time-point as initial conditions, our state-transition models accurately predicted both disease progression and treatment response, supporting this as a potentially valuable approach to time clinical intervention even before phenotypic changes become detectable.

## Introduction

Chronic myeloid leukemia (CML) is characterized at the cytogenetic level by t(9;22), the Philadelphia chromosome that creates BCR::ABL^1^. This fusion gene encodes a ligand-free activated mutant tyrosine kinase that transforms normal hematopoietic stem cells into leukemia stem cells (LSCs), primitive leukemic cells capable of initiating and maintaining the disease.

While the advent of tyrosine kinase inhibitors (TKIs) has revolutionized the treatment of CML by inducing long-term remissions and largely preventing disease evolution from a chronic phase (CP) into blast crisis (BC), these drugs often fail to fully eradicate LSCs^2,3^. Lack of treatment compliance, drug intolerance or acquired BCR::ABL mutations or additional genomic “hits”, therefore may result in disease relapse or progression to BC^4–6^. Thus, while for a subset of patients is possible to eventually discontinue TKIs, most of them remain on lifetime treatment^7,8^.

Quantification of BCR::ABL fusion transcript levels is diagnostic and effective at assessing ongoing treatment response but is less useful for predicting timely disease evolution at the earliest time points. In recent years, the transcriptome has emerged as a promising tool for assessing treatment response and predicting outcome in cancer and leukemia^9,10^. Gene expression profiling using high-throughput technologies has enabled comprehensive analysis of the transcriptome in CML patients and identified gene signatures that reveal molecular mechanisms of transformation and treatment response^11,12^. Recently, Giustacchini et al. reported on a distinct molecular signature of LSCs that selectively persisted after TKI treatment^13^. Ross et al. investigated the transcriptomic landscape of CML patients at diagnosis and during TKI therapy and identified differentially expressed genes and pathways associated with cell proliferation, apoptosis, and drug resistance, thereby predicting treatment response and long-term outcomes^14^. Radich et al. explored transcriptomic changes in CML patients who discontinued TKI therapy and found that specific gene expression patterns, particularly involving immune-related genes, could distinguish patients who successfully maintained treatment-free remission from those who relapsed^15,16^. However, these studies are mainly population-based, and are therefore difficult to apply to predict disease evolution and treatment outcome for individual patients. To our knowledge, no recent studies have examined how transcriptomic changes detected at the earliest time points of disease or at the start of treatment can accurately predict outcome in individual CML mouse or human, even before phenotypic changes had occurred.

Here, we applied state-transition theory to model the temporal dynamics of the CML transcriptome and predict CML evolution and treatment response^9,10^. CML is an ideal model to test this hypothesis as the disease is initiated and initially maintained solely by BCR::ABL, targeted in the clinic by TKIs, a treatment effective in inducing disease remission, but often not curative. We showed that the transcriptome of peripheral blood mononuclear cells (PBMC) from a transgenic CML mouse can be modeled as a particle undergoing state-transition in a CML-transformed potential landscape. Using this model, we accurately predicted the trajectories of disease evolution, treatment response, and relapse of individual mice at the earliest time-point. Furthermore, the state-transition framework allowed for identification of the patterns of gene expression that drove leukemogenesis, thereby connecting transcriptome state-transition to molecular mechanisms of disease growth and treatment resistance.

## Results

### Peripheral blood transcriptome state-transition during CML development

To investigate the molecular mechanisms of CML initiation and evolution, we collected PBMC samples from the CP CML mice at sequential time points after BCR::ABL induction by tetracycline discontinuation (Tet-off; Fig. 1A). Samples were also collected from control mice that continued to receive tetracycline (Tet-on) in order to maintain BCR::ABL suppression. Bulk RNA-sequencing (RNA-seq) was performed on each PBMC sample. To study the transcriptomic changes that characterized disease evolution, we applied a state-transition approach that mapped the time-series transcriptomes of each mouse into a CML state-space. The resulting trajectories of each mouse in the state-space were modeled using a stochastic equation of motion to determine whether the disease evolution of each mouse could be predicted based on its initial condition.

**Figure 1:**
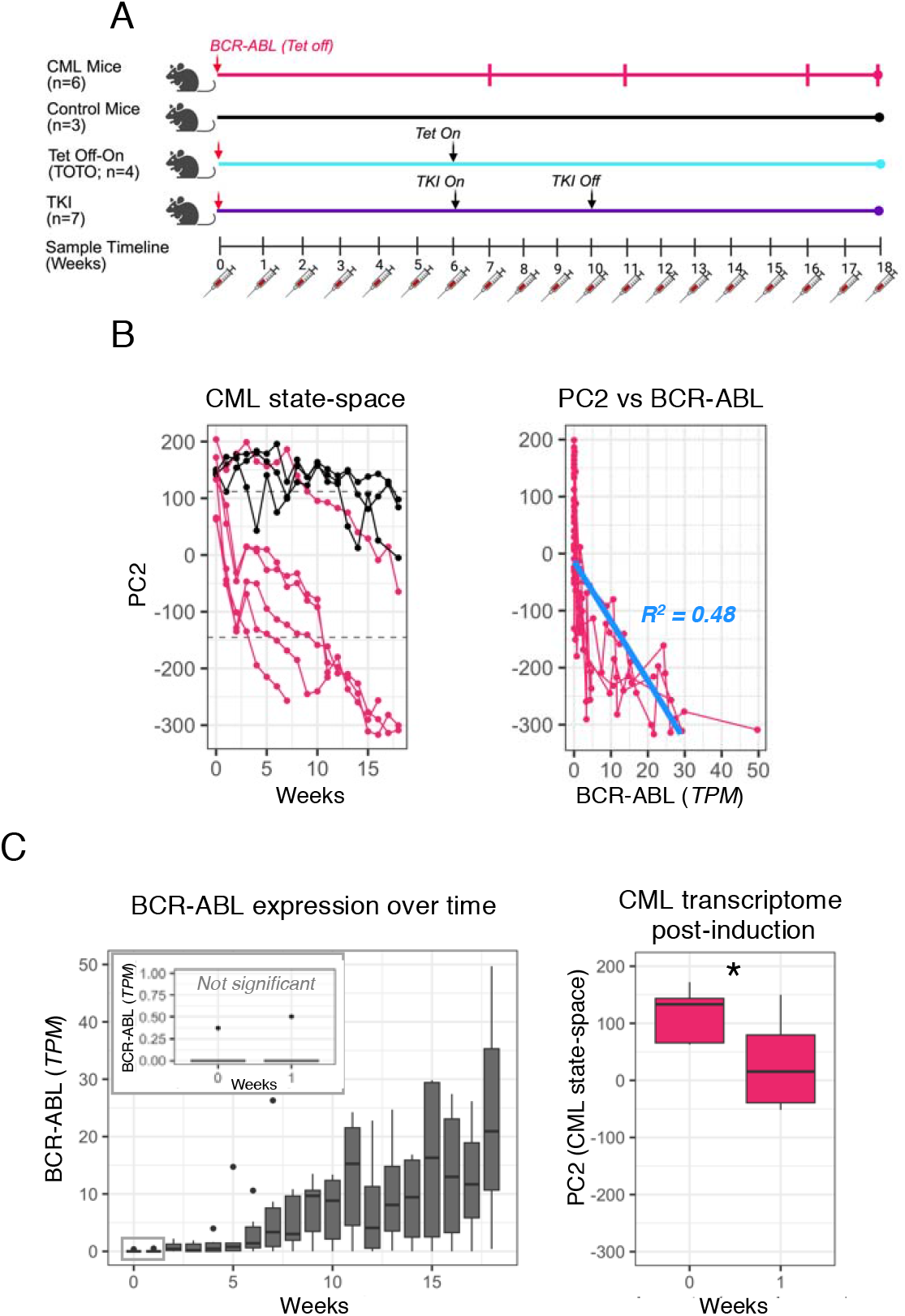
Time-series peripheral blood transcriptome encodes the CML state-space. **A)** The experimental design collected time-series sampling of peripheral blood from four cohorts of mice over 19 weeks. All mice had Tet-inducible expression of the BCR-ABL fusion gene which results in CML when expressed. **B)** Singular value decomposition (SVD) was performed on the full transcriptome using all time points from both the CML and control mouse cohorts to identify the CML state-space (PC2). When plotted against time, the CML state-space shows sample disease trajectories of each sample over time (*left*). PC2 was selected as the CML state-space because it had the largest separation between control mice and the endpoint CML samples, and because it had the best linear fit (in *blue*) with BCR-ABL expression (*right*; Table S1). **C)** BCR- ABL expression only began to increase 4-5 weeks after Tet was turned off (*left*). Whereas BCR- ABL expression remained undetectable at week 1 (*left inset*), the CML state-space already shows a significant movement toward the CML disease state (*right*; p-value=0.006).

To start, the CML state-space was built using Principal Component Analysis (PCA). Briefly, we performed singular value decomposition^9,10^ (SVD; see methods) on all expressed genes (n= 39,927) measured with RNA-seq of the sequentially collected samples of both control and CML mice. By including all the time-sequential samples collected from the CML and control mice, we were able to capture the continuum of transcriptional states that spanned from health to terminal states but also included intermediate states of the disease. Among all PCs, PC2 (hereafter called CML state-space) provided the greatest separation between samples collected from the healthy control mice and those collected from CML moribund mice. We obtained transcriptome trajectories of each individual mouse by plotting the time sequential samples in the CML state-space (Fig. 1B, left panel).

Supporting the ability of the CML state-space to identify distinct disease states, we demonstrated that among all the PCs, PC2 had the best correlation with BCR::ABL expression levels. Nevertheless, the correlation was relatively weak (R^2^ = 0.48; Fig. 1B, right panel; Table S1), suggesting that BCR::ABL was not the only driver of transcriptome state-transition. In fact, immediately after tetracycline discontinuation (Tet-off), we observed that the CML mice began to move through the CML state-space, from the health state toward disease. Despite BCR::ABL expression not yet being detectable, it likely influenced disease state changes even at those lowest, initial levels (Fig. 1C).

### CML transcriptome state-transition model predicts disease evolution

As the CML state-space contained the continuum of the transcriptomic states from health to disease, we used the state-space to build a state-transition model that predicted the disease trajectories of individual mice. These predictions were made by setting initial conditions for each mouse using only the state-space location from the first collected time point. The theoretical framework of this model postulated that changes in the transcriptome could be modeled as a particle undergoing Brownian motion in a potential energy landscape using the Langevin equation^9,10^. Thus, the solution to the Fokker-Planck (FP) equation corresponding to the equation of motion was used to predict disease state-transition in the CML state-space.

To understand the system dynamics, we hypothesized that the CML potential energy landscape could be modeled using a three-node network that resulted in a tri-stable system (Fig. 2A)^17^. In this system, BCR::ABL was the input signal that interacted with the PBMC cell population. The changing PBMCs and the consequent transcriptome variation created a double negative feedback loop representing how leukemogenesis dysregulated the transcriptome and how the dysregulated altered transcriptome affected the composition (normal vs leukemic) of the cell population. The transcriptome, represented by the location of the sample in the CML statespace, was modeled as self-activating progressive dysregulation of gene networks during leukemia progression^18^. The potential landscape resulting from this system ranges from a single well centered at the health state (BCR::ABL off) to three wells representing the most energetically favorable, CML steady states (BCR::ABL maximum; Fig. 2B). The equations describing the dynamic of the transcriptome in the CML potential resulting from this three-node network are described in the Methods.

**Figure 2:**
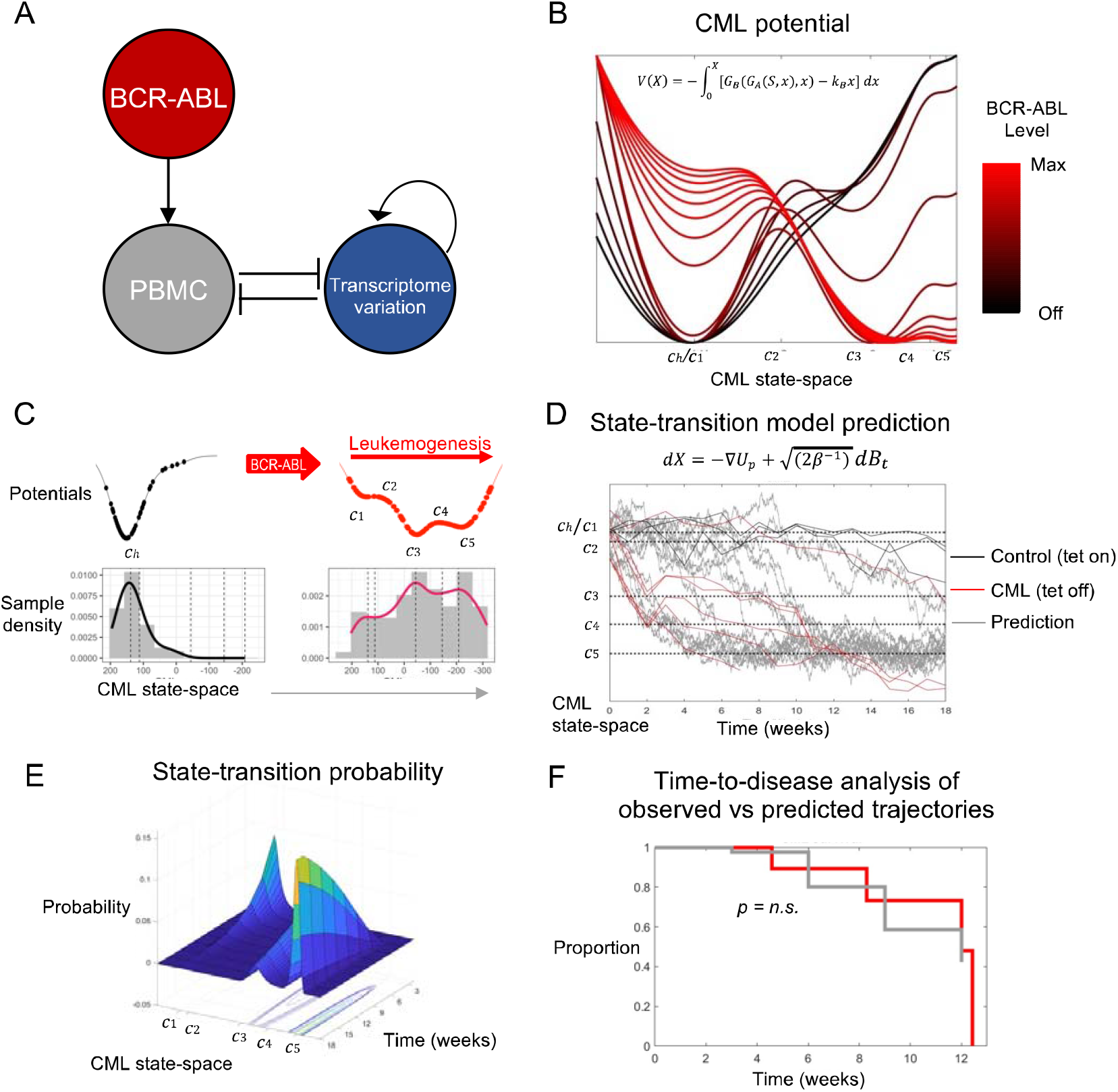
State-transition model accurately predicts time-to-disease. **A)** The three node CML network model that produced a tri-stable systems and captures the features of the transcriptomic CML potential. **B)** BCR-ABL input signal determines the shape of the CML potential. When BCR-ABL is off (*black*), the system has one steady state at the healthy (*c*_*h*_) state. As the BCR-ABL signal increases, the potential gains additional steady states. When BCR-ABL signal reaches its maximum, which we use to model CML development in this experiment, the potential has five critical point which we labeled *c*_1_ *= c*_5_. Three of these critical points are steady states (*c*_1,_ *c*_3,_ *c*_5_), and the most energetically favorable state (*c*_5_) is the CML disease state. **C)** Using the distribution of samples in the CML state-space, we empirically constructed the control (*left*) and CML (*right*) potentials. CML potentials were created by fitting histograms of samples in the CML state-space with kernel densities (bottom). The polynomials that defined potentials (*top*) were then constructed by finding the critical points of the density curves and using those as the zeros of the derivative of the potential. **D)** Using only the first (week 0) time point of each sample as an initial condition, the stochastic equation of motion was solved forward in time for the CML and control potentials to predict the trajectory of each sample in the CML state-space (*grey lines*). **E)** The solution of the state-transition model is given as a probability density over time which represents the likelihood of finding a sample in the CML state-space as time evolves. **F)** To test the accuracy on the state-transition model predictions, a time-to-disease analysis was performed showing that there was no significant difference between the predicted and observed trajectories (log-rank p-value = 0.8) and that they were highly similar (concordance index = .75).

To test if the theoretically-derived leukemogenic potential reliably recapitulated the experimental observations, we considered the empirical distribution of the CML mouse samples in the statespace and derived an experimental CML potential (Fig. 2C; see Methods). The tri-stable CML potential was characterized by five critical points identified from the kernel density estimator of the empirical distribution (denoted *c*_1_–*c*_5_). We labeled the stable (minima) critical points as Early state (Es) at *c*_1_, Transition state (Ts) at *c*_3_, and Late state (Ls) at *c*_5_ (Fig. 2C). The unstable (maxima) critical points were labeled as Early-Transition state (T-Es) at *c*_2_ and Transition-Late state (T-Ls) at *c*_4_ ; these unstable points represent the boundaries between two stable states. We also used the sample density of the control mice to determine the location of a Health state (Hs) labeled *c*_*h*_ as the single stationary state and critical point for the control mice. The samples from the control mice remained confined to a region of the CML state-space corresponding to the Health state (Hs) *c*_*h*_. Importantly, the three wells of the experimentally derived potential directly mapped into those of the theoretically predicted potential (Fig. 2B-C).

To predict time-to-disease [i.e., the amount of time that elapsed before the mouse crossed the boundary of the Ls (*c*_5_)], we modeled the location and trajectory of each mouse’s transcriptome in the state-space using the CML potential and the Langevin equation of motion. We were able to generate predicted transcriptome trajectories that matched the observed ones using only the data from samples collected at the first time point (Fig. 2D). The Fokker-Planck (FP) solution of the Langevin equation, when solved forward in time, accurately predicted time-to-disease (Fig. 2E). As all the CML mice died after passing Ls (*c*_5_), we also compared the time-to-disease (defined as the first crossing of the Ls (*c*_5_) critical point) between the observed and the predicted trajectories. The predicted and observed time-to-disease curves overlapped and were not statistically different (p = 0.8, Log-rank; Fig. 2F), and were highly concordant (C-index = 0.75) demonstrating accuracy of the state-transition model’s prediction. Of note, one CML mouse never developed disease after Tet-off and remained healthy (Fig. 2D). Our model predicted that this mouse would not cross Ls (*c*_5_), thereby accurately reproducing the experimental observation.

### State-transition critical points of BCR::ABL potential landscape defined distinct disease states

In order to identify the transcriptomic drivers of CML state-transition, we reasoned that samples from different mice must be compared at equivalent disease states. Since mice developed CML at different times, we could not simply rely on grouping transcriptome samples by their collection time point. Thus, we utilized the samples’ positions in the state-space to group them into distinct disease states. We then compared gene expression at each disease critical point disease [i.e., Es (*c*_1_), Ts (*c*_3_) and Ls (*c*_5_)] with that of control samples at health critical point [i.e., Hs (*c*_*h*_) ; Fig. 3A; Fig. S3A; Table S2].

**Figure 3:**
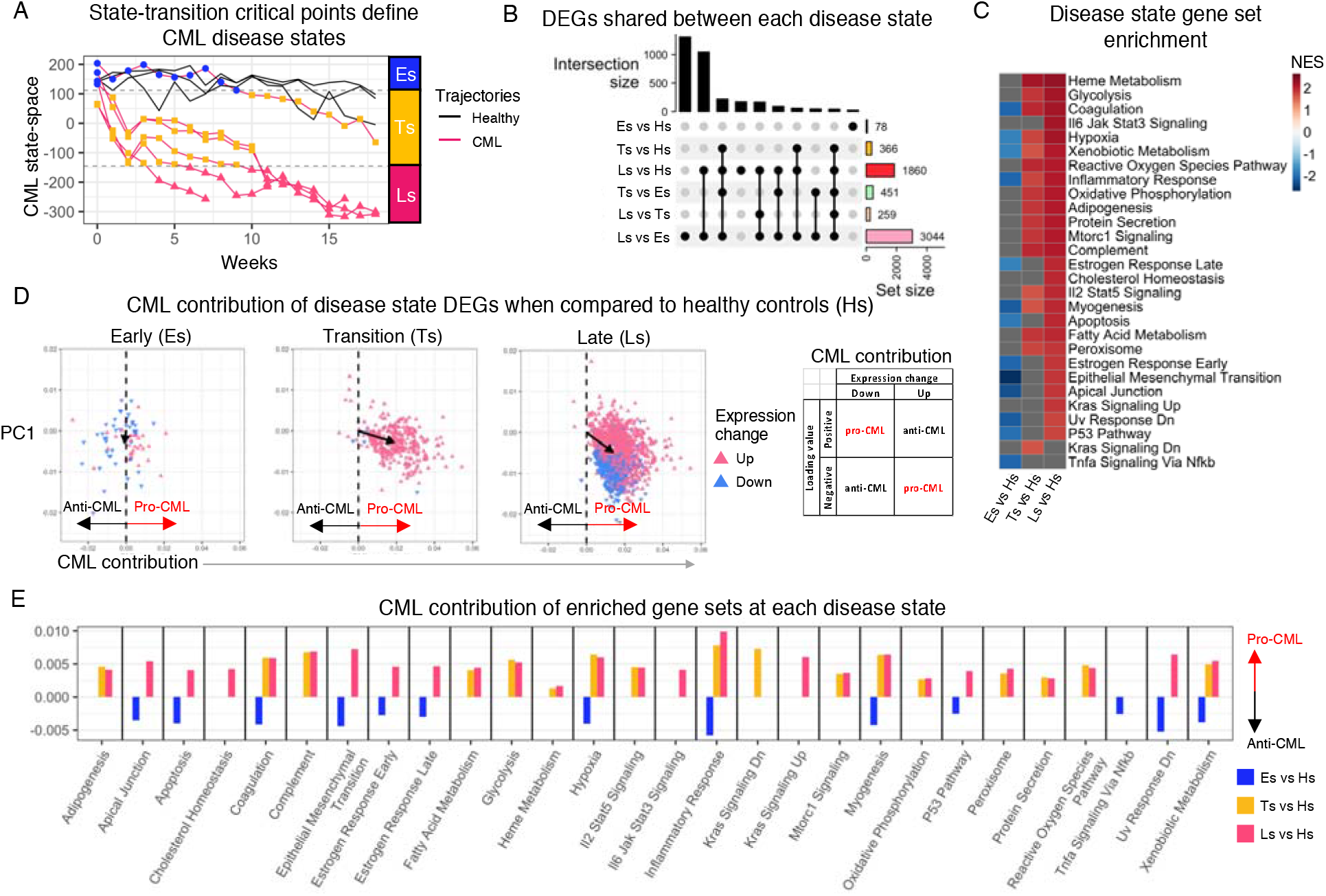
Differential expression analysis performed on CML state-space disease states. **A)** The three stable critical points of the CML state-space (*c*_1,_ *c*_3,_ *c*_5_) were used to group the CML samples into Early (Es), Transition (Ts), and Late (Ls) disease states based on their location in the CML state-space. The unstable critical points (*c*_2_, *c*_4_; *dashed lines*) were used as the boundaries to define the three disease states. **B)** Differential gene expression was performed both between the disease states and healthy control state (Hs) and between the disease states. The number of differentially expressed genes (DEGs) contained in the intersection between the comparisons was used to determine how similar the disease states were to each other. **C)** Gene set enrichment analysis was performed on the log_2_ fold change of the healthy vs disease states to determine which Hallmark gene sets were significantly enriched (between disease states shown in Fig. S3B). The results were summarized by showing a heatmap of the normalized enrichment score (NES) for all significant pathways (non-significant pathways are in *grey*) across the three disease states. The NES score shows the direction of expression change for each gene set. **D)** Using the eigengenes (PC2 loading values) used to construct the CML states-space, we determined the contribution to CML of each gene that was a DEG in the disease state vs control comparisons. The CML contribution is determined by combining the eigengene value with the observed expression change for that comparison (*right box*). The total contribution for each DEG comparison was indicated by the mean vector (*black arrow*) of all DEGs in each comparison. **E)** To visualize the contribution of each significantly enriched gene set, the total CML contribution (mean vector) for each comparison was plotted for each comparison (missing bars indicate non-significant gene set in that comparison).

A total of 78 differentially expressed genes (DEGs) were identified at Es (*c*_1_) as compared to Hs (*c*_*h*_). A large proportion of DEGs (55 out of 78 total DEGs) observed at the Es (*c*_1_) state were unique to this state and were not identified in the other disease states (Fig. 3B). Among the most upregulated genes, were Ins1 and Ins2 that encode insulin, in addition to Fam177a1 and several SnRNAs (Table S2). Fam177a1 encodes a protein localized to the Golgi complex and endoplasmic reticulum but of unknown function, and when mutated, is associated with developmental and neurological disorders^19^. Of note, BCR::ABL was not among the upregulated DEGs at the Es (*c*_1_). Among the most downregulated DEGs, there were several genes encoding for collagen and elastin. Compared to Hs (*c*_*h*_), the transition state Ts (*c*_3_) presented with 366 DEGs (Fig. 3B; Table S2). In addition to BCR::ABL, among the most upregulated DEGs, was Prrt4, encoding for a protein predicted to be an integral component of the cell membrane, and the lncRNA Gm6093, predicted to play a role in cell differentiation^20,21^. Among the most downregulated DEGs, we observed collagen and elastin encoding genes, in addition to Il33. Compared to Hs (*c*_*h*_), Ls (*c*_5_) presented with 1860 DEGs (Fig. 3B; Table S2). In addition to BCR::ABL, among the most upregulated DEGs, there were Pgk1-rs7, involved in glycolysis, and Prrt4. Among the most downregulated DEG, there were several pseudogenes and other genes coding proteins with unknown function. Of note, 94% of DEGs identified at Ts (*c*_3_) were also DEGs and expressed at higher levels at Ls (*c*_5_) (Fig. 3B).

Using gene set enrichment analysis (GSEA), we identified Hallmark gene sets enriched at distinct disease states (Fig. 3C). Compared with Hs (*c*_*h*_), we observed that all significantly enriched gene sets were downregulated at Es (*c*_1_). These included Epithelial Mesenchymal Transition (EMT), Apical Junction, and Myogenesis, that have been previously associated with malignant transformation^22–24^. In contrast, compared with Hs (*c*_*h*_), all significantly enriched gene sets were upregulated in the Ts (*c*_3_) and Ls (*c*_5_) (Fig. 3C). Among the most upregulated gene sets, we found those related to metabolism, angiogenesis, and inflammation. Of note, while all enriched gene sets at the Ts (*c*_3_) and Ls (*c*_5_) were upregulated with respect to Hs (*c*_*h*_), we observed that these genes sets had higher expression at Ls (*c*_5_) compared with Es (*c*_1_), with the exception of Pancreas Beta Cells which was downregulated (Fig. S3B).

### Eigengene analysis quantifies DEG contribution to CML development

To quantify the contribution of each DEG to CML development, we used the CML state-space loading value of each gene which we defined as CML eigengenes^25^. Located in the second column of the matrix *v*^*^ from the SVD procedure that was used to construct the CML statespace (Table S2), the eigengene value described the direction and the magnitude of the effect that each gene had on the sample’s location in the state-space. Thus, by combining the eigengenes with the expression changes at each disease critical point with respect to Hs (*c*_*h*_), we quantified the impact of DEGs on disease onset and evolution and categorized each gene as either a pro-or anti-CML eigengene. A pro-CML eigengene had either a positive loading value with downregulated expression or a negative loading value with upregulated expression; an anti-CML eigengene had either a positive loading value with upregulated expression or a negative loading value with downregulated expression (Fig. 3D; S4A). The eigengenes were then visualized in a 2D space constructed by plotting the CML contribution vs the PC1 loading value (Fig. S4A). Of note, since PC1 was not directly associated with BCR::ABL expression or CML development, its selection was made simply to map the CML contribution; any other PC could have been used instead.

The net contribution to CML of the Es (c_1_), Ts (*c*_3_) or Ls (*c*_5_) DEGs was calculated as the vector mean of the vectors representing individual DEGs (Fig. 3D; Fig. S4B). The DEGs at Es (*c*_1_) had a net anti-CML contribution, while those at Ts (*c*_3_) and Ls (*c*_5_) had a net pro-CML contribution.

We also calculated the CML contribution of the significantly enriched gene sets as a mean vector of all the eigengenes in a given enriched gene set. All the Es (*c*_1_) enriched gene sets had a net anti-CML contribution (n=13), whereas all the Ts (*c*_3_) and Ls (*c*_5_) enriched gene sets had a net pro-CML contribution (n=17 and 26 respectively; Fig. 3E; Fig. S4C).

To assess whether the eigengene-based contributions corresponded to phenotypic differences between disease states, we compared myeloid counts at Es (*c*1), Ts (*c*3), and Ls (*c*_5_) states. We observed no myeloid cell growth at Es (*c*_1_); this was followed by an initial expansion at Ts (*c*_3_) and further growth at Ls (*c*_5_) (Fig. S3D). The growth of the myeloid population between Ts (*c*_3_), and Ls (*c*_5_) was associated with an expansion of the DEGs at Ls (*c*_5_) and an increase in enrichment in many of the shared processes (Fig. 3B; Fig. S3B). GSEA indicated these genes to be members of metabolism- and inflammation-related gene sets. In addition, 405 downregulated DEGs were unique to the Ls (*c*_5_) state (Fig. S3C; Table S2). However, over half of the downregulated DEGs were unnamed genes and failed to be significantly enriched in any pathway (265 of 405). Nevertheless, the eigengene analysis showed that all but one of the downregulated Ls (*c*_5_) DEGs had a pro-CML contribution.

### Gene expression dynamics identify drivers of CML state-transition

While we used the state-space critical points as a classifier of distinct disease states, the continuous state-space coordinates could also be used to align transcriptomes by their progress toward CML. By doing so, we created a pseudotime ordering of the disease, which allowed us to identify the patterns of gene expression involved in CML transformation. To this end, we used a correlation-based analysis to identify subsets of genes with similar patterns of expression during disease evolution (hereafter called gene “modules”; Fig. S5). At Es (*c*_1_), we found no gene modules, meaning that the DEGs identified at Es (*c*_1_) were not co-regulated at subsequent states (Fig. 4A, left; Fig. S5A). At Ts (*c*_3_), we identified an “increasing module”, meaning that these DEGs were co-regulated and continued to increase during disease progression (Fig. 4A, middle; Fig. S5B). At Ls (*c*_5_), we identified two gene “modules”, one increasing and one decreasing (Fig. 4A, right; Fig. S5C).

**Figure 4:**
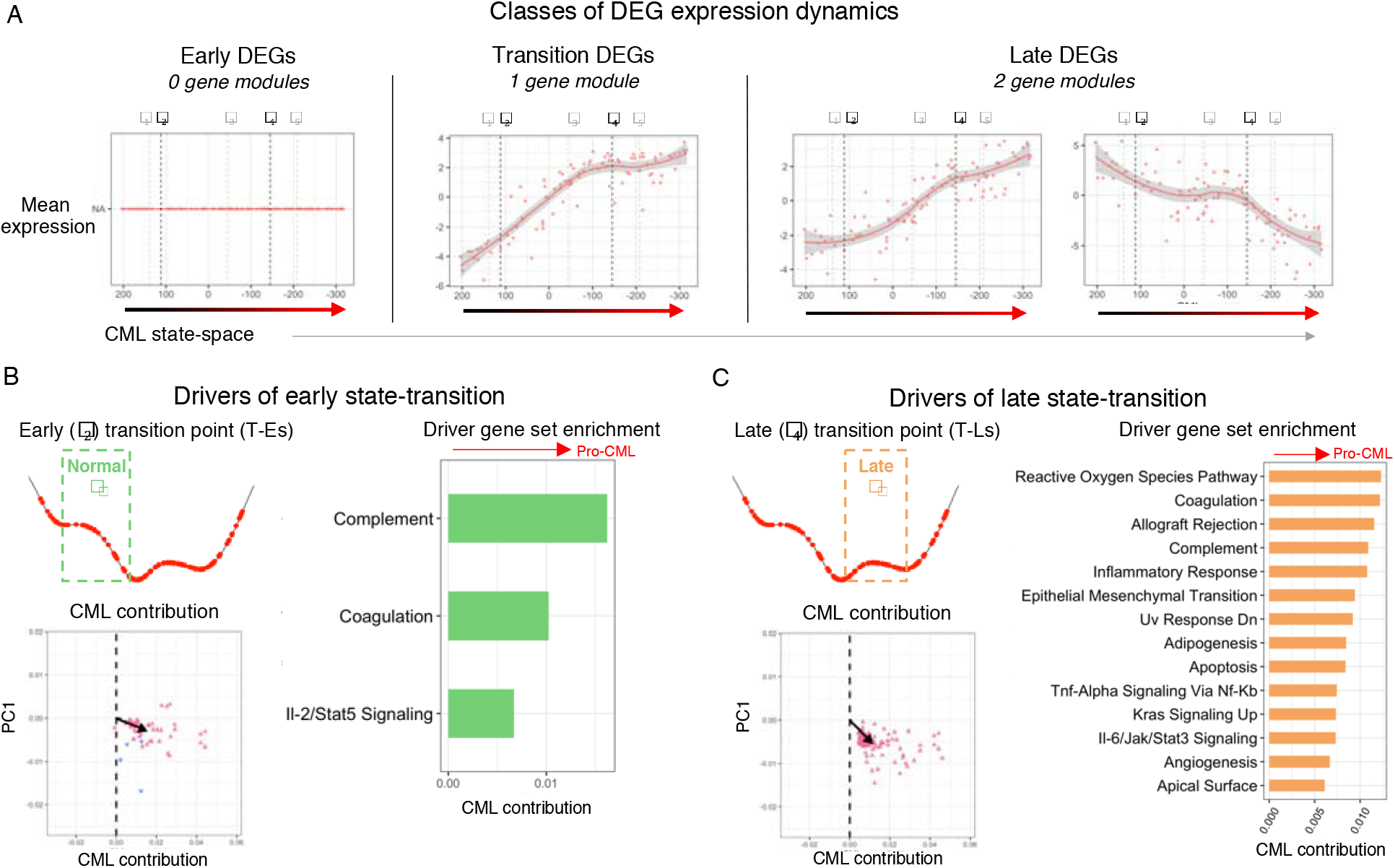
CML state-space aligns samples and identifies transition driver genes. **A)** Using a correlation analysis to identify the DEGs with similar dynamics during CML development, gene modules were identified for the unique DEGs of the three disease states (Es, Ts, Ls; Fig. S3C, S4A). To visualize the expression of each detected gene module that had a mean correlation coefficient > 0.25, the mean log-normalized mean-centered expression of all genes in the module was plotted as function of the CML state-space coordinate for each CML sample. **B-C)** For the early transition point (T-Es; **B**) and the late transition point (T-Ls; **C**) DEGs that were involved in state-transition were identified by correlating the expression of each DEG with the shape of the potential around each of the transition points (*left top*; Fig. S6*B*). To identify the driver gene processes, the driver genes for each transition point were then extended to their high-confidence interaction partners using STRINGdb (Fig. S6C). The CML contribution were summarized for the resulting protein-protein interaction networks (*left bottom*). The processes enriched by the STRINGdb extended driver genes involved were identified and their CML contribution was calculated (*right*).

Different from the increasing gene module observed at Ts (*c*_3_), the increasing gene module at Ls (*c*_5_) had non-linear dynamics, as it presented an inflection point near T-Ls (*c*_4_) (Fig. 4A). The inflection represented the point where the rapid increase of DEG expression started to slow down and became constant. The “decreasing” Ls (*c*_5_) gene module also had a non-linear expression curve, with an inflection occurring at approximately T-Ls (*c*_4_) and continued through Ls (*c*_5_) (Fig. 4A). It is important to emphasize that the expression dynamics of the gene modules were not derived from any critical point information; rather were derived independently, using only the mean expression of the DEGs and the CML state-space location of each transcriptome sample. Thus, it was striking to us that the inflections of the gene modules occurred almost exactly at T-Ls (*c*_4_), an unstable critical point between Ts (*c*_3_), and Ls (*c*_5_). The observation of co-regulation at the unstable critical points led us to hypothesize that these types of coordinated expression changes could break the equilibrium of a stable critical point and drive the disease transition from that stable state of the disease to the next one.

To identify the driver genes of disease transition between two otherwise stable states, we correlated the expression of individual DEGs with the shape of the CML potential (Fig. 4B-C; Fig. S6A). For this analysis, we included all DEGs (n=3,304) resulting from the comparisons of the stable disease states with the health state (Hs). We identified 194 DEGs at the T-Es (*c*_2_) state and 497 at the T-Ls (*c*_4_) state whose expression followed the CML potential (Fig. S6B). All but one of the T-Es (*c*_2_) genes and all T-Ls (*c*_4_) genes had a pro-CML contribution, which supports the hypothesis that these genes drove the CML state-transition. Interestingly, only 23 DEGs were identified as being involved in both T-Es (*c*_2_) and T-Ls (*c*_4_) transitions, suggesting that unique and distinct processes likely drove disease evolution from across the distinct disease states (Fig. S6B).

To investigate which cellular processes were involved in the transition between two stable states [i.e., Es (*c*_1_) to Ts (*c*_3_) and Ts (*c*_3_) to Ls (*c*_5_)], we constructed protein-protein interaction networks and performed pathway enrichment analysis (Fig. S6C). Both T-Es (*c*_2_) and T-Ls (*c*_4_) were significantly enriched for protein interactions (p<0.0001; 31 and 232 interacting proteins respectively). The T-Es (*c*_2_) transition point was characterized by inflammation and immune related processes and involved multiple collagen genes (Fig. 4B; Table S3). T-Ls (*c*_4_) transition point was characterized by metabolism-related processes in addition to inflammation- and immune-related processes also enriched in T-Es (*c*_2_) (Fig. 4C, S6C; Table S3). All enriched gene sets for both T-Es (*c*_2_) and T-Ls (*c*_4_) had a pro-CML contribution, supporting their role in driving disease evolution.

### State-space transition predicts treatment response

To investigate how treatment impacted CML state-transition, we performed time-series experiments using two approaches (Fig. 1A, 5A, S7A). The first approach made use of the CML mouse model’s ability to induce and eventually suppress BCR::ABL expression upon tetracycline discontinuation (Tet-off) and subsequent re-administration (Tet-on); hence we referred to this treatment as Tet-off-Tet-on (TOTO)]. The TOTO approach represented a hypothetical best scenario where a treatment is able to completely suppress BCR::ABL expression (Fig. S7A). The second approach consisted of treatment with nilotinib, a TKI used in clinical practice (hereafter referred to as “TKI”). TKI was assumed to be less effective than TOTO as, rather than suppressing BCR::ABL expression, it inhibited BCR::ABL at the protein level. Both Tet-on and TKI treatments were initiated six weeks after Tet-off. In the TOTO group, Tet-on was continued for 12-weeks. In the TKI cohort, TKI was administered for 4 weeks, and the mice were followed for the subsequent 12 weeks, for a total of 18-weeks or until they succumbed to disease, whichever occurred first.

To determine the effect of each therapy on CML state-transition, the transcriptome dynamics of mice receiving one of the two treatments were compared with those of the untreated diseased and healthy mice at each of the disease [(Es (*c*_1_), Ts (*c*_3_), Ls (*c*_5_)] or health Hs (*c*_*h*_) states respectively; Fig. 5B, S7B; Table S4). In order to assess post-treatment CML disease states, we used the eigengenes to project the TOTO and TKI samples into the CML state-space (see Methods). Supported by an observed relationship between the number of DEGs and the mean location of sample groups in the state-space, we used both the number of DEGs and the statespace to assess transcriptomic similarities or differences (Fig. S7C). We defined treatment response as movement of the transcriptome in the state-space away from the CML LS (*c*_5_) state and toward Hs (*c*_*h*_), with a complete response being a return to health [Hs (*c*_*h*_)].

**Figure 5:**
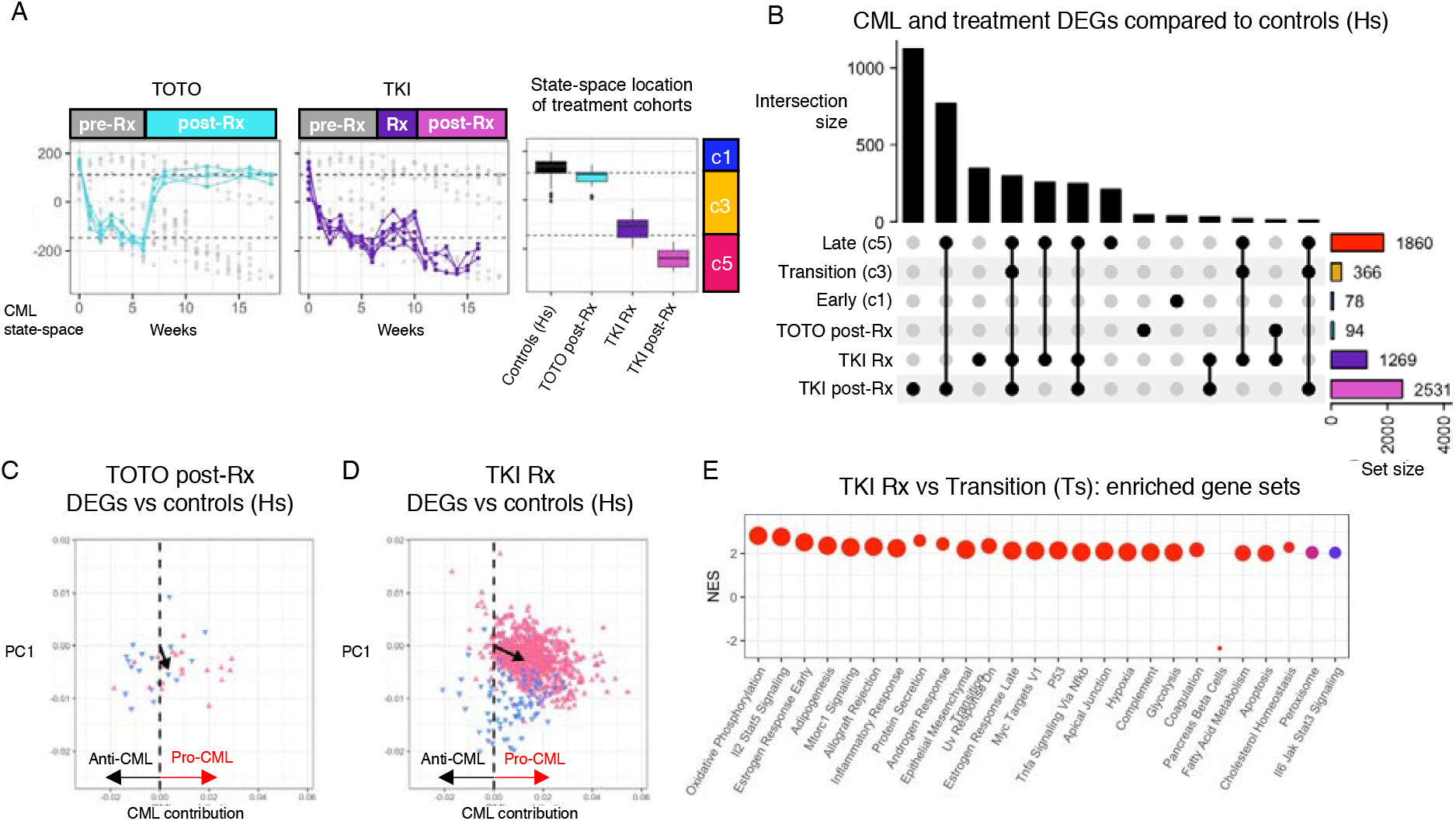
CML treatment effects determined using CML state-space disease states. **A)** Samples from the TOTO (*left*) and TKI (center) were projected into the CML state-space to allow the treatment effects to be compared to the CML disease states (*right*). **B)** DEGs were identified by comparing the TOTO post-Rx and both the TKI Rx and post-Rx samples to the controls (Hs). All intersections with 10 or more DEGs are shown (all intersections shown in Fig. S7B). **C)** After removing the age-related DEGs that were detected when comparing the early vs late control samples, the CML contribution of the 38 TOTO post-Rx vs controls DEGs were plotted and showed a small overall pro-CML contribution (*black arrow*). **D)** The CML contribution of each of the TKI Rx vs control (Hs) DEGs are shown along with the overall CML contribution for all DEGs (*black arrow*). **E)** Since the TKI Rx samples returned to a similar CML state-space location, the significantly enriched (adjusted p-value < 0.0001) GSEA Hallmark gene sets for the TKI Rx vs transition (Ts) DEGs were determined to identify what processes were affected by TKI treatment.

When disease states of TOTO and TKI treated mice were compared with those of controls, we observed significantly different results. In the TOTO cohort, we observed a rapid movement from LS (*c*_5_) toward Hs (*c*_*h*_) in the state-space soon after tetracycline was re-administered. Posttreatment TOTO samples were tightly distributed around the *c*_2_ critical point, which represented a near-health, newly created stable state in the treatment transformed potential landscape (Fig. 5A). When treated mice at *c*_2_ were compared to the controls, few DEGs were identified (Fig. 5A- B). The observed DEGs and enriched gene sets that were detected by comparing TOTO mice to healthy controls were similar to those observed as time-related effect in the control mice alone and therefore, were possibly associated with aging (Fig. S7D). However, after subtracting out the time-related DEGs, 38 DEGs remained and had a small pro-CML contribution (Fig. 5C), indicating that once BCR::ABL transformation had occurred, the transcriptome maintained some leukemic fingerprints even if BCR::ABL expression is completely suppressed.

Different from TOTO, TKI treatment moved the mouse transcriptomes more slowly from the initial pre-treatment Ls (*c*_5_) state. These transcriptomes reverted only to Ts (*c*_3_) and none of these mice achieved the response of the TOTO mice (Fig. 5A). When compared to controls, the TKI treated mice had 1269 DEGs (Fig. 5B), and a net pro-CML contribution with BCR::ABL the top pro-CML DEG (Fig. 5D). Using GSEA to compare TKI treatment with healthy controls, gene sets that were upregulated during CML remained upregulated during TKI treatment (Fig. 5E) and had an overall pro-CML contribution (Fig. S7F). Of note, BCR::ABL was the top ranked pro- CML DEG.

### Modeling the effects of TOTO and TKI on the BCR::ABL potential

We next tested whether treatment could alter the potential landscape and, in turn, whether the state-transition model could predict treatment response at the earliest time point after treatment initiation. We started by identifying which parameters should be perturbed to recapitulate the state-transition observed with each of the treatments. To model TOTO treatment, we set the

BCR::ABL signal to zero(*S* = 0) and introduced a treatment response parameter *γ*_*TOTO*_ which modulated the action of the signal on the transcriptome as 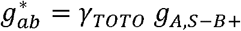 with *γ*_*TOTO*_ =7.5 (Fig. 6A left). This resulted in a potential with a new steady state occurring near the T-Es (*c*_2_) state, the exact location in the state-space where the TOTO mice mapped post-treatment (Fig 6A right). We observed that both a decrease in BCR::ABL expression and an increase in the transcriptome sensitivity to BCR::ABL were necessary to derive a post-treatment potential that matched the observed TOTO trajectories. Once BCR::ABL signal was set to zero, the modulation of the single parameter 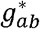 was sufficient to transform a pre-treatment tri-stable CML potential(*V*_*CML*_*(x)*) into a system with only one stable state corresponding to *c*_2_ following TOTO treatment (*V*_*TOTO*_ (*x*)). Using the TOTO potential (*V*_*TOTO*_ (*x*)), we then produced simulated TOTO trajectories that matched those observed (Fig. 6B; Fig. S8).

**Figure 6:**
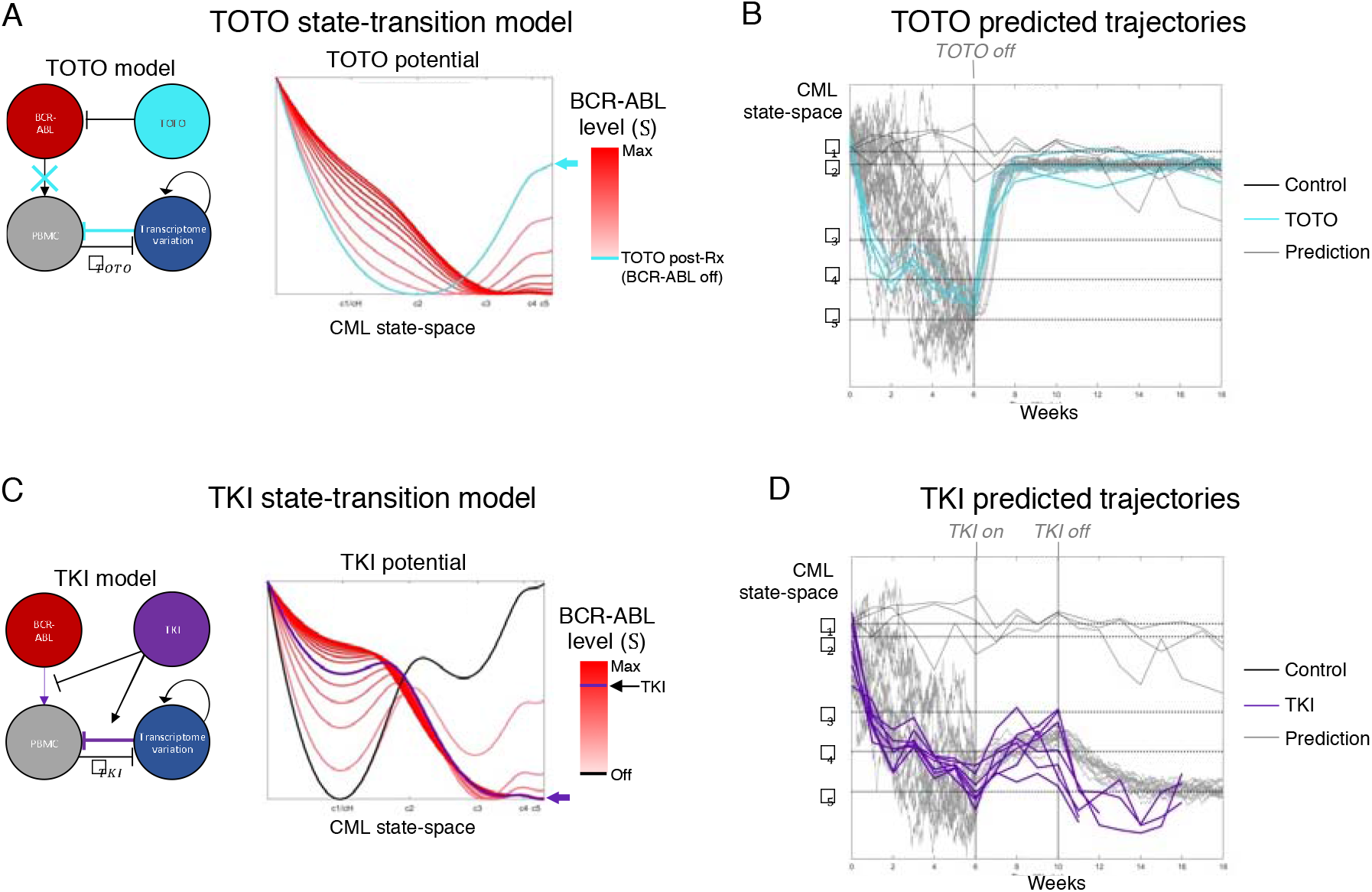
State-transition model predicts CML treatment dynamics. **A)** The model for the TOTO cohort is modified from the CML model to include turning off the BCR-ABL input signal and increasing the *γ*_*TOTO*_ term in the model (*left*). As the input BCR-ABL signal is decreased in this model, the potential transforms from a tri-stable CML potential (*red curve*) to a potential with one state that has minimum energy at the unstable transition critical point c_2_ (*teal curve*; *right*). **B)** Using the TOTO potential, sample trajectories were predicted by solving the stochastic equation of motion forward in time using the first time point as an initial condition. **C)** The model for the TKI cohort was modified to reduce the BCR-ABL signal and increase the *γ*_*TKI*_ term in the model (*left*). The TKI potential (*purple*) lies between the CML (BCR-ABL max) and the BCR-ABL off potentials. The model showed that the healthy state (Hs) could only be achieved as an energetically favorable state while on TKI was for the BCR-ABL signal approach zero. **D)** Using the TKI potential, sample trajectories were predicted by solving the stochastic equation of motion forward in time using the first time point as an initial condition.

To model TKI treatment, we reduced the BCR::ABL signal (BCR::ABL > 0) and introduced a treatment response parameter *γ*_TKI_ (Fig. 6C left). In contrast to TOTO, the post TKI potential was produced only when both the BCR::ABL signal was reduced (*S*< *S*_max_) and the response rate between the PBMCs and the BCR::ABL signal was increased, 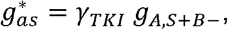, with *γ*_TKI_ =1.4. The change in this single parameter was sufficient to transform the tri-stable CML potential (*V*_*CML*_ (*x*)) into a bi-stable potential with new stable states at *c*_1_ and *c*_3_ following TKI treatment (*V*_TKI_(*x*)). Importantly, the TKI model predicted that the only way TKI treated mice could return to Hs (*c*_*h*_) was if the BCR::ABL level approached zero (Fig. 6C right). When we used this potential to simulate TKI treatment, we again observed an accurate prediction of the observed transcriptome trajectories (Fig. 6D; Fig. S8).

## DISCUSSION

We have previously reported that state-transition models can be used to predict time-sequential transcriptome dynamics and disease evolution in biological systems such as murine acute myeloid leukemia (AML)^9,10,18^. Constructing a state-space to model biological transitions can capture changes produced by genetic, epigenetic, or microenvironmental perturbations on the transcriptome, which simultaneously deregulate multiple biological processes in order to generate a phenotype such as the leukemic transformation observed here^26–29^. We have previously shown that micro-RNA and protein abundance (proteomic) data can also be used to create a state-space and characterize state-transitions^10,30^. Using a transgenic CP CML mouse model that recapitulated the human disease, we report here the application of state-transition theory to interpret the temporal transcriptomic changes involved both in CML development and in treatment (where CML is “reversed”) to accurately predict outcome and treatment response.

We demonstrate the feasibility of predicting state-transition dynamics and time to disease using the time-sequential transcriptomic data collected from a murine model of CML. Our results show that the movement of the blood transcriptome in the state-space could be modeled as that of a particle undergoing Brownian motion in a leukemogenic potential defined by stable and unstable critical points. We hypothesized that BCR::ABL creates a leukemogenic potential that arises due to the action of BCR::ABL on both the PBMCs and the transcriptome. We also derived a CML potential landscape based on the position of the critical points derived from the observed distribution of the CML mouse transcriptomes in the state-space. The theoretically and experimentally derived landscape potentials were qualitatively identical and could be used to predict the trajectory of disease development over time. We showed that at the earliest time points, the CML state-space was more sensitive at detecting the disease state transition than the BCR::ABL expression itself. Furthermore, we showed that our state-transition model predicted transcriptome trajectories that matched well the observed ones utilizing only the transcriptomic state derived from data collected at the first time point after Tef-off BCR::ABL induction.

The biological features that characterized the distinct disease states were inferred from the analysis of DEGs obtained by comparing gene expression at each of the CML critical points with the control health state. Because the SVD decomposed the gene expression into linear combinations of orthogonal basis vectors, we were able to assess the contribution of individual genes, DEGs, or entire Hallmark gene sets to CML initiation and growth. Although the SVD can produce artifacts when applied to time-series data, we observed no oscillations or artifacts, and moreover, we observed a strong correlation between state-space trajectories and BCR::ABL expression as an immunophenotypic marker of disease, supporting our interpretation of the principal component as a transcriptome state-space^31^. We observed that the Es (*c*_1_) state was largely characterized by anti-CML DEGs that resisted CML development whereas the Ts (*c*_3_) and Ls (*c*_5_) disease states were characterized by pro-CML DEGs associated with the movement of state-transition toward late disease state. In other words, the anti-CML contributions present only at Es (*c*_1_), were suggestive that at this early state of the disease, the transcriptome changes opposed BCR::ABL-driven disease development, whereas the pro-CML contribution as observed at Ts (*c*_3_) and subsequently at Ls (*c*_15_) suggested that transcriptome changes at these points overwhelmed any initial anti-leukemic force and contribute to disease evolution.

Interestingly, while we noted that genes involved in inflammation and angiogenesis were enriched at both Ts (*c*_3_) and Ls (*c*_5_), we also observed upregulation of genes involved both in glycolysis and oxidative metabolisms at Ls (*c*_5_), suggesting an increase in the bioenergetic demand in the state of highest growth^5,32,33^. Studying the changes in the expression of individual DEGs, we also discovered that groups of co-regulated genes (i.e., gene modules) that increased or decreased concurrently. Some of these gene modules displayed non-linear expression trends with inflection points that mirrored the shape of the CML potential at the unstable critical points Es (*c*_2_) and Ts (*c*_4_). Therefore, we hypothesized that they were likely the major drivers of transition between two stable disease states. Among others, metabolic processes were involved in the transition between Ts (*c*_3_) and Ls (*c*_5_), suggesting that the increased metabolic demand was one of the possible causes of the transition to the final state of the disease, when maximum leukemic growth was observed.

Importantly, we also applied the state-transition model to predict response to therapy. To this end, first we treated CML mice in Ls (*c*_5_) with TOTO, which resulted in complete suppression of BCR::ABL. We showed that TOTO brought the system rapidly back close to Es (*c*_1_), without reaching it. In fact, TOTO-treated mice remained at a new stable critical point TOTO (*c*_2_), with a small number of pro-CML DEGs. Thus, it is likely that once the system is altered, some fingerprints of BCR::ABL transformation remain, despite complete suppression of the fusion gene. Importantly, using only the initial time point and treatment start date, our potential model accurately predicted treatment response and post-treatment dynamics.

In contrast, TKI treatment returned the transcriptome only to Ts (*c*_3_). Once TKI treatment was completed, the transcriptome rapidly returned to Ls (*c*_5_) and mice had disease relapse. This mimics a possible trend observed in those patients that are either not compliant to the treatment and need to stop due to intolerable side-effects or have developed TKI resistant disease that evolves to a blast phase. While the model also accurately predicted TKI trajectories, it also predicted that with administration of higher dose or more effective TKIs, the system may potentially return to the healthy state, Hs (*c*_*h*_). However, the optimal dosage of TKI compatible with the return to health may not be attainable in vivo^34^. Although the treatment response parameters between TOTO and TKI may not be directly compared, it is interesting that the treatment response effect for TOTO (*γ*_*TOTO*_) was higher than that for TKI (*γ*_*TKI*_) suggesting that the suppression of BCR::ABL gene with tetracycline had a greater positive impact on the transcriptome than BCR::ABL protein inhibition by TKI.

In summary, we show here that state-transition dynamical models can predict disease evolution and treatment response in a murine model that recapitulates human disease. It is possible therefore that this approach could be applied to other -omic data types and translated into the clinic as an assay to predict disease evolution and treatment response at the earliest time points, even before phenotypic changes will occur.

## METHODS

### Mouse model

Inducible transgenic SCLtTA/BCR::ABL mice of B6 background were maintained on tetracycline (tet) water at 0.5 g/L. Tet withdrawal (tet-off) results in expression of BCR::ABL and generation of a CML-like disease in these mice with a median survival of approximately 10 weeks after induction of BCR::ABL, which recapitulates human CP CML. Six to eight weeks old male and female SCLtTA/BCR::ABL mice were randomly divided into 4 groups: control (Tet on, n=3 mice), CML (Tet off, n=6 mice), Tet Off-On (TOTO; Tet off 6 weeks then Tet on for 12 weeks, n=4 mice), TKI [Tet off 6 weeks then TKI (Nilotinib, 50mg/kg, oral gavage, daily) treatment 4 weeks, n=7 mice]. Blood was collected from these mice weekly, i.e., before BCR::ABL induction (t = 0) and weekly after induction up to 18 weeks (t = 1 to 18) or when the mouse developed leukemia and became moribund, whichever event occurred first.

### RNA-seq sequencing, data processing and analysis

For each mouse, peripheral blood samples were accrued for all timepoints before for RNA extraction. Total RNA was extracted from whole blood using the AllPrep DNA/RNA Mini Kit (Qiagen, Hilden, Germany); quality and quantity were estimated using a BioAnalyser (Agilent, Santa Clara, CA). Samples with a RIN > 4.0 were included. Sequencing libraries were constructed using the KapaHyper with RiboErase (Kapa Biosystems, Wilmington, MA), loaded on to a cBot system for cluster generation, and sequenced on a Novaseq 6000 (Illumina) with paired end 100-bp for mRNA-seq to a nominal depth of 40 million reads. To mitigate batch effects, samples were assigned to flow cells such that treatments were approximately evenly represented across runs and samples from the same mouse shared a run. Image processing and base calling were conducted using Illumina’s Real-Time Analysis pipeline.

Raw sequencing reads were processed with the nf-core RNASeq pipeline version 3.7^35^. Briefly, trimmed reads were mapped using STAR to the GRCm39 reference (Genbank accession GCA_000001635.9) amended with the human *BCR::ABL1* fusion gene sequence (Genbank accession EF158045.1) using GENCODE annotation Release m28^36^. Each library was subjected to extensive quality control, including estimation of library complexity, gene body coverage, and duplication rates, among other metrics detailed in the pipeline repository^35^. Transcript abundance was estimated across genomic features using Salmon and merged into a matrix of pseudocounts per million transcripts (TPM) per gene for each sample^37^. *BCR::ABL1* transcript abundance was measured by counting reads that spanned the fusion boundary. Surrogate variable analysis was used to check for confounding experimental effects^38^. None was apparent (data not shown); however, for all comparisons, sex was used as a covariate in differential expression analyses using the default setting of DESeq2 and counts generated by tximport using the Salmon abundance estimation^39^. The RNA-seq dataset is submitted to the Gene Expression Omnibus and assigned accession number GSE244990.

### State-space construction

To construct the CML state-space, we performed singular value decomposition (SVD) on all expressed genes in the transcriptome (n=39,927), which we defined as all genes expressed in at least one sample. By including all time points of the time-series peripheral blood samples in both the control and the BCR::ABL expressing CML mice, we both captured the full transcriptional system and spanned all intermediate disease states (Fig. 1B). The SVD is a dimensionality reduction technique that produces low rank approximations of the input data, called principal components (PCs), that capture sources of variation in the original data set (Fig. S1B). Among the principal components, we found an axis on the PC1 vs PC2 plane to be the component with the best separation between the control and CML endpoint samples. To align this axis with the y-axis, we found the rotation angle that made the linear fit line of all control samples parallel to the x-axis. After applying a 37.8° rotation to the PC1 vs PC2 plane, we identified the rotated PC2 to be the principal component that best encoded CML development. We determined this based on two criteria: 1) it had the largest separation between the control samples and the endpoint samples for the CML mice; and 2) it demonstrated the best correlation with both BCR::ABL expression (abundance and log-normalized) and the myeloid population in the CML mice (Fig. S1A; Table S1). Therefore, we defined PC2 as the CML statespace. When we plotted PC2 against time for all samples, the CML state-space provided disease trajectories for each mouse (Fig. 1B). Although the correlation with BCR::ABL was by far the best for the CML state-space compared to all other PCs, the correlation was low (R^2^ = 0.48; Table S1). Although PCA can be misinterpreted when applied to time-series data with temporal correlations, we are confident in our application and interpretation here because of the correlation with BCR::ABL expression, phenotypic disease manifestation, and the absence of oscillations or other artifacts in the PCs^31,40^. This is due to the orthogonality of the control (tet on) and CML (tet off) mice transcriptome endpoints and dynamics, which enable an orthogonal decomposition of the data and identification of the leukemia signal from the background/noise.

Using the expression of the same genes used when performing SVD to construct the statespace, the projection of the TOTO and TKI treatment cohorts was performed as follows: the treatment expression values *X*_T_, were multiplied by the state-space loading values,*<u>V</u>*, and the inverse of the eigenvalues,∑^-1^ (Fig. 3B). The second column of the resulting treatment cohort PC matrix,U_*T*_, represented the coordinates of the treatment samples in the CML state-space.

### Mathematical model

The three compartment mechanistic model used to generate the CML, TOTO, and TKI potentials is adapted from Dey and Barik 2021.^17^ Using the double negative feedback loop B (DNFL-B) circuit framework, BCR::ABL concentration is modeled as the signal (S) acting on PBMC cells (A) and the corresponding observable variation in the transcriptome (B), as

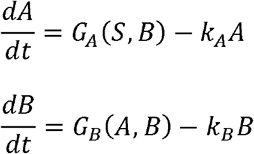

where *G*_*A*_ (*S,B*)and *G*_*B*_(*A,B*) are rate functions relating the concentration of the signal to the cell and transcriptional states as

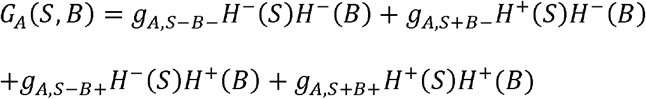

and similarly for *G*_*B*_ (*A,B*), see Dey and Barik 2021. The functions *H*^-^ and *H*^+^ are switching Hill functions given by

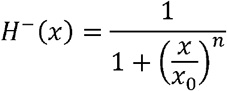

and *H*^+^ (*x*) = 1− *H*^−^(*x*).The parameters are as stated in Dey and Barik Table S2 for the DNFL- B circuit model.

The effective potential *V* (*x*)for the system is given by integrating the effective force of the signal (S) on the cells (A) and observable transcriptome (B) as

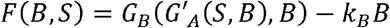

Where G^′^_A_is the steady state rate of 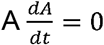, so that

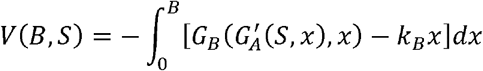

is a function of the state variable *B*, which in our case is the state-space coordinate corresponding to PC2, which parametrically depends on the signal, or BCR::ABL expression level *S*.

### Mean squared displacement analysis

Mean squared displacement (MSD) was calculated for each mouse from their CML state-space trajectories vs time (Fig. S2A). The MSDs of each experimental group was fit to a linear model and the slope of the line was used to estimate the diffusion coefficient (*β*) used in the state-transition model (Fig. S2B).

The MSD at a time *t* is given as *MSD* =⟨|*x*(*t*)−*x*_*m*_|^2^⟩, where *x*(*t*) is the position of trajectory from the simulation by a treatment model and *x*_*m*_ is the CML state-space trajectory of the mouse model. The MSD was calculated for each mouse for each time point *t*, and the average MSD for all time points was evaluated by changing the coefficient of treatment force *γ* and post-treatment diffusion coefficients *β*. The MSDs were examined within the range of *γ* for 0.1 ≤ *γ* ≤ 2.0 and *β* For0.001≤ *β* ≤0.1, and the combination of values of *γ* and *β* that has the minimum MSD was selected. The search ranges of parameter, *β*, was empirically estimated from the average slope of linear regression to CML and control mouse data. The upper bound of *γ* was determined by the convergence of trajectories of the simulation.

### DEG expression dynamics

Expression dynamics were investigated by performing hierarchical clustering on the correlation matrixes produced using all CML mouse time points for each of the unique Early, Transition, and Late DEGs. Genes with similar expression dynamics (gene modules) were determined by selecting cutting the dendrogram to obtain clusters and then selecting for clusters that had a median correlation coefficient greater than 0.25 (Fig. S5A-C). For each gene module, the expression dynamic plots were created by calculating the mean of the log-normalized mean-centered expression for each CML sample and then plotting that value vs the CML state-space coordinate of the sample (Fig. 4A).

### Transition point driver genes

To identify which genes had expression dynamics that were most similar to the inflection points of the CML potential, we used the following approach (Fig. S6A). First, we included all DEGs obtained by comparing CML disease state vs both healthy (*c*_*h*_) and other CML disease states in the analysis. Second, we defined each transition point by selected the CML samples that were located near the T-Es (*c*_2_) (CML state-space coordinates = [*c*_1,_ *c*_3_]) or the T-Ls (*c*_4_) (CML statespace coordinates = [*c*_3,_ *c*_5_]). Finally, we performed both a correlation analysis and linear regression separately for the T-Es (*c*_2_) and the T-Ls (*c*_4_) transition points between the lognormalized mean-centered expression and the value of the potential at the corresponding CML state-space of each sample. The driver genes for each transition point were those genes with a correlation coefficient greater than 0.5 and a linear regression significance of p < 0.05 (Fig. S6B). To identify the processes that were involved at each transition point, we used STRINGdb protein-protein interaction database to extend the driver genes to all high-confident (interaction score > 900) interaction partners (Fig. 4B-C).^41^ Using the drivers plus their interaction partners, EnrichR gene set enrichment analysis was performed on the Hallmark gene sets to identify the significantly enriched transition point processes (Fig. 4B-C).^42,43^ Since the T-Ls network was quite large and included a number of enriched gene sets, we identified the comprising subnetworks and performed EnrichR on each subnetwork to better resolve the involved processes (Fig. S6C)

### TOTO and TKI treatment simulations

We integrated treatment into the state-transition treatment model by modeling the treatment as a force that affects both the potential 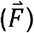 and internal energy of the particle, or the diffusion rate (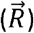). The equation of motion for the transcriptome particle is then given by the Langevin equation as

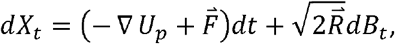

where the position of the particle in the CML potential is denoted by *X*_*t*_, and *β*_*t*_ is a Brownian stochastic process that is uncorrelated in time ⟨*B*_*i*,_ *B*_*j*_ ⟩ = δ_i,j_.

The corresponding probability distribution *p*(*x,t*) at time *t* is given by a Fokker-Plank (FP) equation including the treatment force 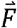 and diffusion vector 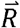 as follows:

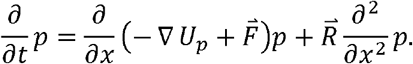

We described the Tet-off CML potential as ∇*U*_*p*_ *=a*(*x*− *c*_1_) (*x*− *c*_2_) (*x*− *c*_3_) (*x*− *c*_4_) (*x*− *c*_5_), with critical points *c*_1_, *c*_2_, *c*_3_, *c*_4_ and *c*_5_ as identified in the state-space. The state-transition potential *U*_p_ was mapped to the potential *V*_*CML*_ (*x*)using the critical points in the state-space as follows.

For TOTO treatment, the CML and normal state-transition potentials approximated those generated by the network model as

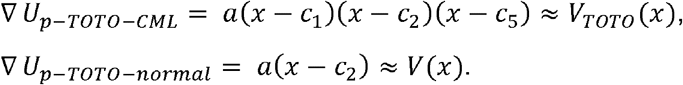

The transformation from CML to normal was therefore given as

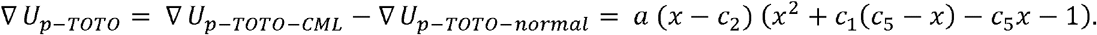

The TOTO treatment started by reintroducing Tet at *t*=6 until the study endpoint, therefore, the treatment force, 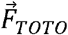, and diffusion vector 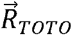 were given as follows:

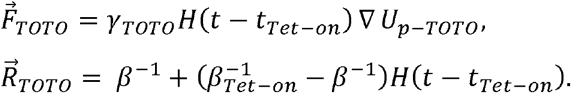

The parameter *γ*_*TOTO*_ represents the treatment strength, *H* is the Heaviside step function and *t*_*Tet*-*on*_ is the timing when Tet treatment started.

For TKI, the CML and normal potentials were approximated using critical points and midpoints between *c*_1_ and *c*_5_ based on the observations from the network model given as follows:

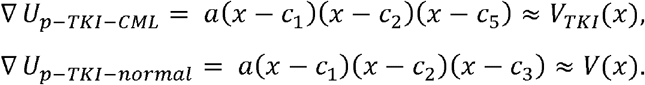

The transformation from CML to normal for TKI treatment was therefore given as

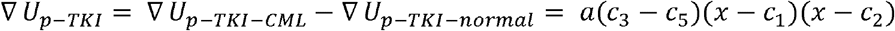

The TKI treatment was introduced from *t*=6-9, incorporating the nilotinib half-life, *γ*.^44^ The treatment force for TKI,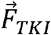, and 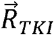, were defined as the following,

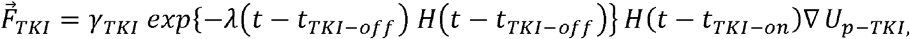

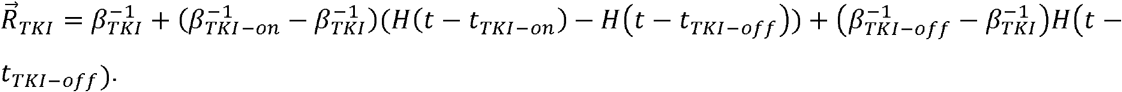

In the model, the transition from a normal potential to a CML potential was approximated using three critical points using *c*_1_, *c*_2_ and *c*_5_ as ∇*U*_p-*CML*_ *=a*(*x*− *c*_1_) (*x*− *c*_2_) (*x*− *c*_5_), and∇*U*_p-*normal*_ *=a*(*x*− *c*_1_) (*x*− *c*_5_).

A FP equation was solved to generate the probability density in the state-space for each treatment, TOTO, TKI, and CML using the state-transition potentials. The FP equation for the TOTO system is given by

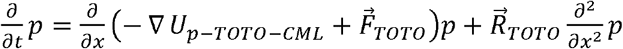

For TKI, the FP equation is given by

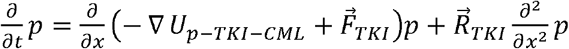

and

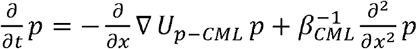

for the CML system. The probability density and trajectories at a given time were quantified by solving each FP solution with the initial condition given by the data at the initial time point.

**Table 1.**
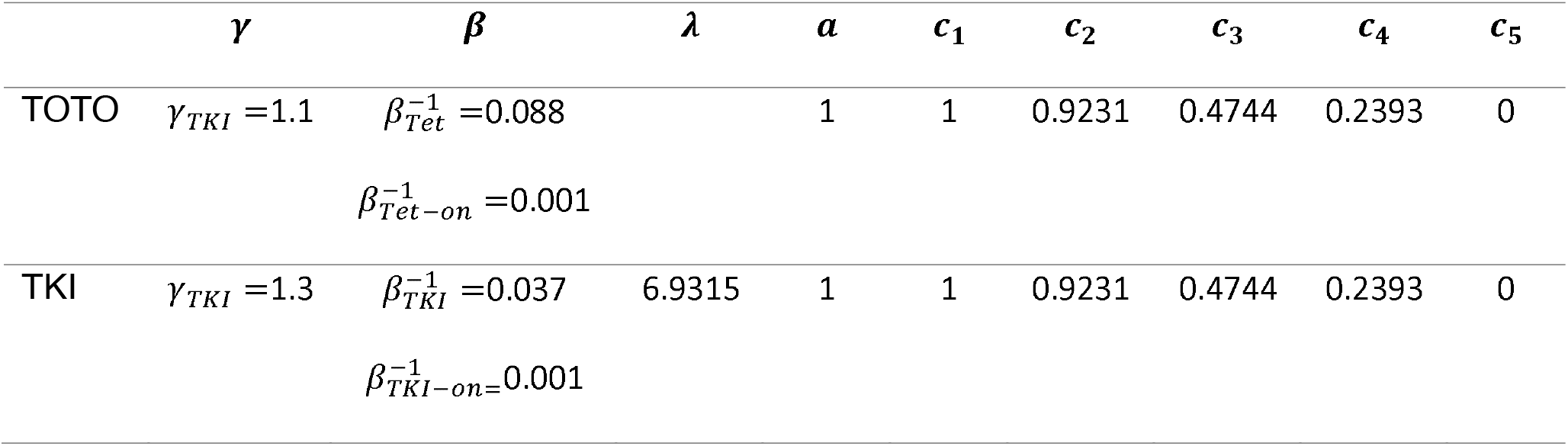
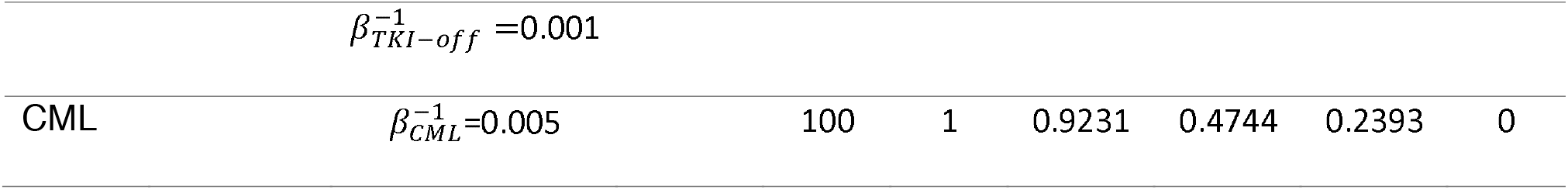
Table 1 Values of parameters used in the modeling.

### Time to disease prediction

To compute the probability of disease progression, we integrated the solution of the Fokker-Planck equation beyond the critical point *c*_5_ over time as

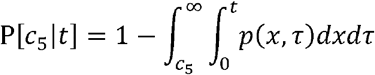

where *p*(*x,t*) is the solution to 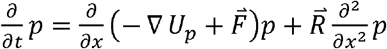 with initial conditions given by the PC2 values at time t=0 of the CML (tet-off) samples. The continuous curve given by P[*c*_5_ |*t*] was then evaluated at the sample observation timepoints, corresponding to 1-week intervals. The observed time to disease was given by the timepoint at which the CML state-space trajectory crossed the *c*_5_ critical point and was computed for each mouse. The predicted and observed time to disease curves were compared using the log-rank test (p=0.88). As an additional statistical test, we applied the concordance index (C-index = 0.747) to show that the predicted time to state-transition was concordant with the observed^45,46^.

### Early vs late control analysis

An analysis comparing the early (first five time points) vs late (last five time points) control samples was performed to determine how the samples changed over time since they were observed to move downward in the CML state-space (Fig. S7D; *top*). The total CML contribution of the 118 DEGs identified showed a small pro-CML contribution in the late controls (Fig. S7E; *bottom left*). The significantly enriched Hallmark gene sets from GSEA were very similar to those identified when the TOTO post-Rx were compared to all Hs samples (Fig. S7E; *bottom right*). Because of the small observed pro-CML effects observed in this analysis that could be due to aging processes in the mice over the course of the experiment, the TOTO post-Rx DEGs were controlled for aging by removing the DEGs observed from the early vs late control analysis.

## Code availability

All code used to analyze and model the data and to generate figures and tables are available at: https://github.com/cohmathonc/CML_mRNA_state-transition

## Acknowledgement

Research reported in this publication was supported in part by Robert and Lynda Altman Family Foundation and included work performed in the integrative genomics core and biostatistics and mathematical oncology Shared Resource supported by the National Cancer Institute of the National Institutes of Health under grant number P30CA033572 and grant U01CA250067. The content is solely the responsibility of the authors and does not necessarily represent the official views of the National Institutes of Health.

## Supplemental Figures and Table

**Figure S1:**
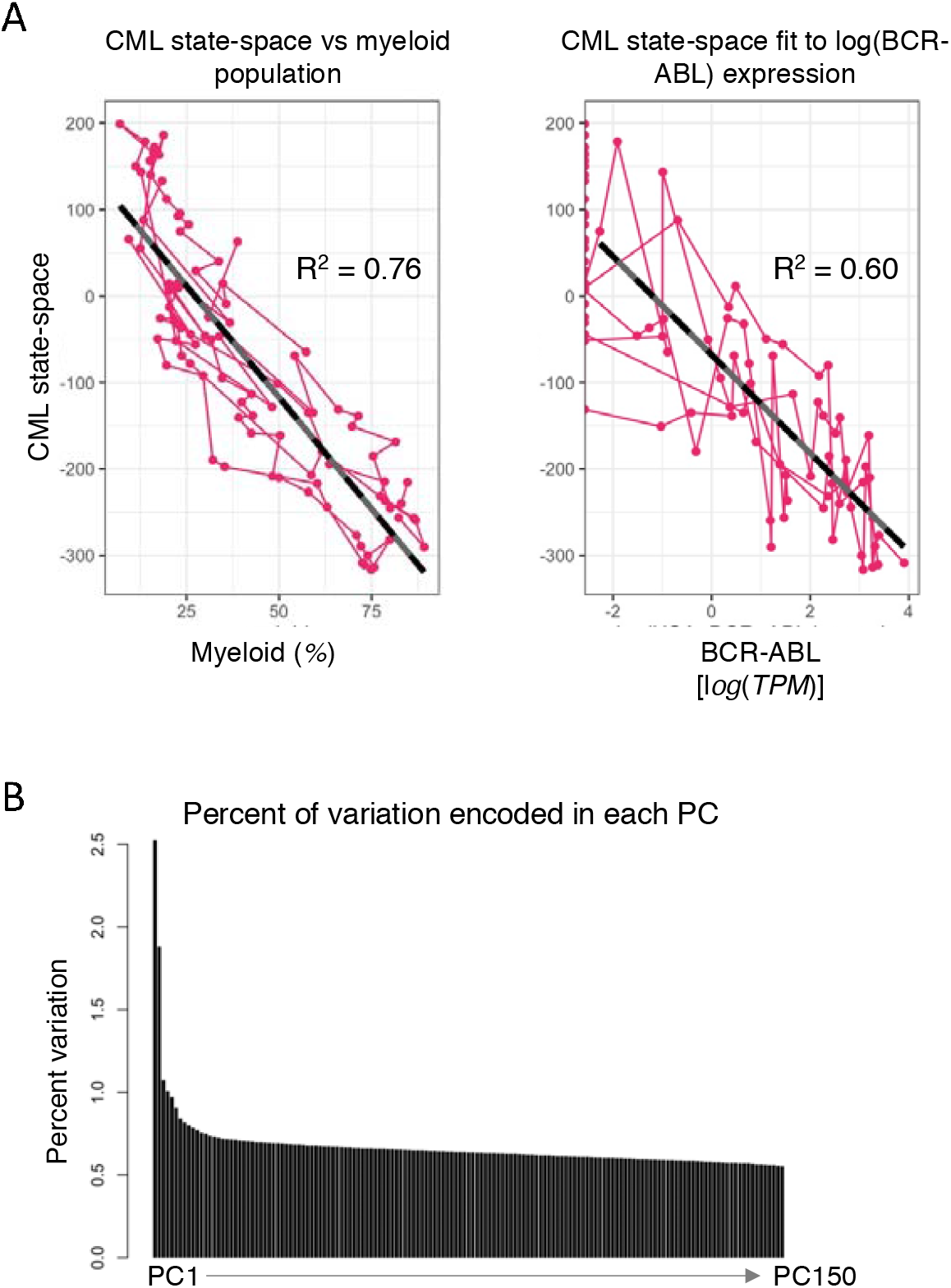
Features of the CML state-space. **A)** The CML state-space was correlated with clinical markers of CML the myeloid population in the peripheral blood (*left*) and the log-expression of BCR-ABL (*right*). Linear fits (*black dashed*) were used to show that the CML state-space had the best correlation with both myeloid and BCR-ABL (Table S1). **B)** The percent of variation encoded by each PC that resulted from the SVD operation was plotted and showed that the variation encoded by PCs decreased and leveled off after PC2

**Figure S2:**
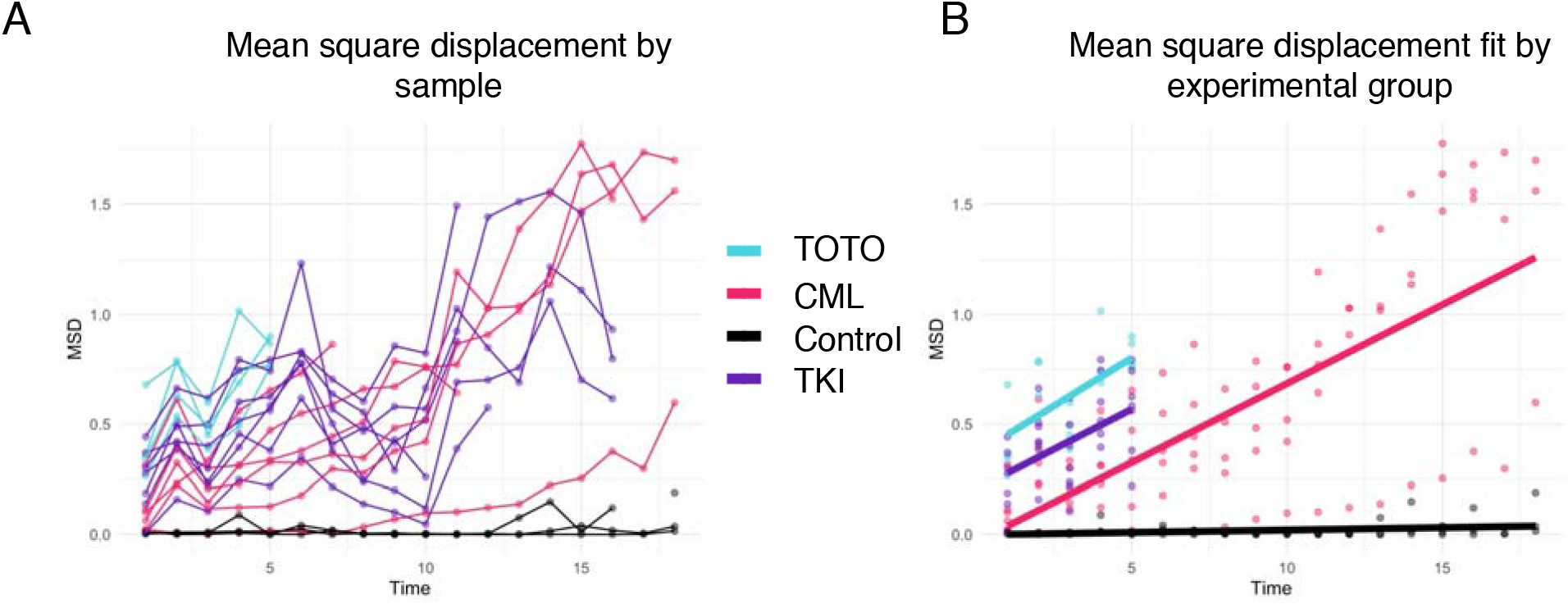
Mean squared displacement estimates for the diffusion coefficients. **A)** A mean square displacement (MSD) was performed for each mouse using their trajectories in the CML state-space as a function of time. **B)** For each cohort of mice, a linear fit of all trajectories in the cohort was calculated and the slope of the line was used to estimate the diffusion coefficient (β). Treatment cohorts were shown only until treatment was initiated.

**Figure S3:**
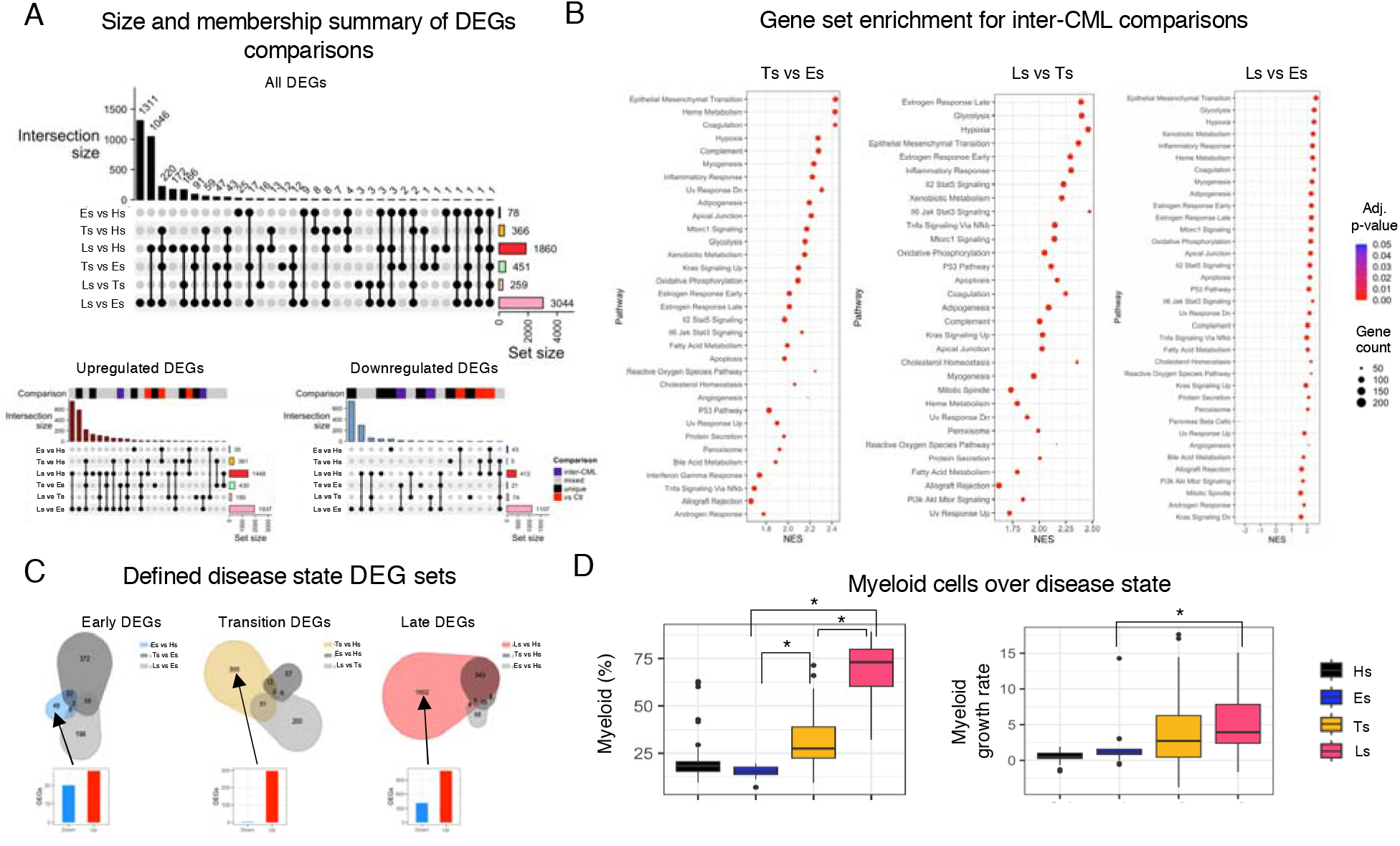
Analysis of CML disease states. **A)** Differentially expressed genes (DEGs) were identified between all pairwise comparisons of the CML disease states (Es, Ts, Ls) and healthy controls (Hs). The results for the intersection between each comparison were plotted separately for all genes (*top*), the upregulated genes in each comparison (*left*), and the downregulated genes in each comparison (*right*). **B)** Gene set enrichment analysis (GSEA) was also performed on all comparisons using the Hallmark gene sets. Here, the inter-CML disease state comparisons (pairwise comparisons of Es, Ts, and Ls) were plotted using the normalized enrichment score (NES) to show the direction of the expression in all significantly enriched gene sets (adjusted p-value < 0.001). **C)** The DEGs unique to each disease state were defined by making intersections between the relevant comparisons to determine which gene expression changes only occurred in each disease state. **D)** The percent of myeloid cells in the peripheral blood was compared in each disease state using a t-test (*left*). The myeloid growth rate was calculated as the derivative of the spline fits for each mouse’s myeloid percentage (*right*) and a t-test was again used to compare growth rate between disease states (significant p-value < 0.05).

**Figure S4:**
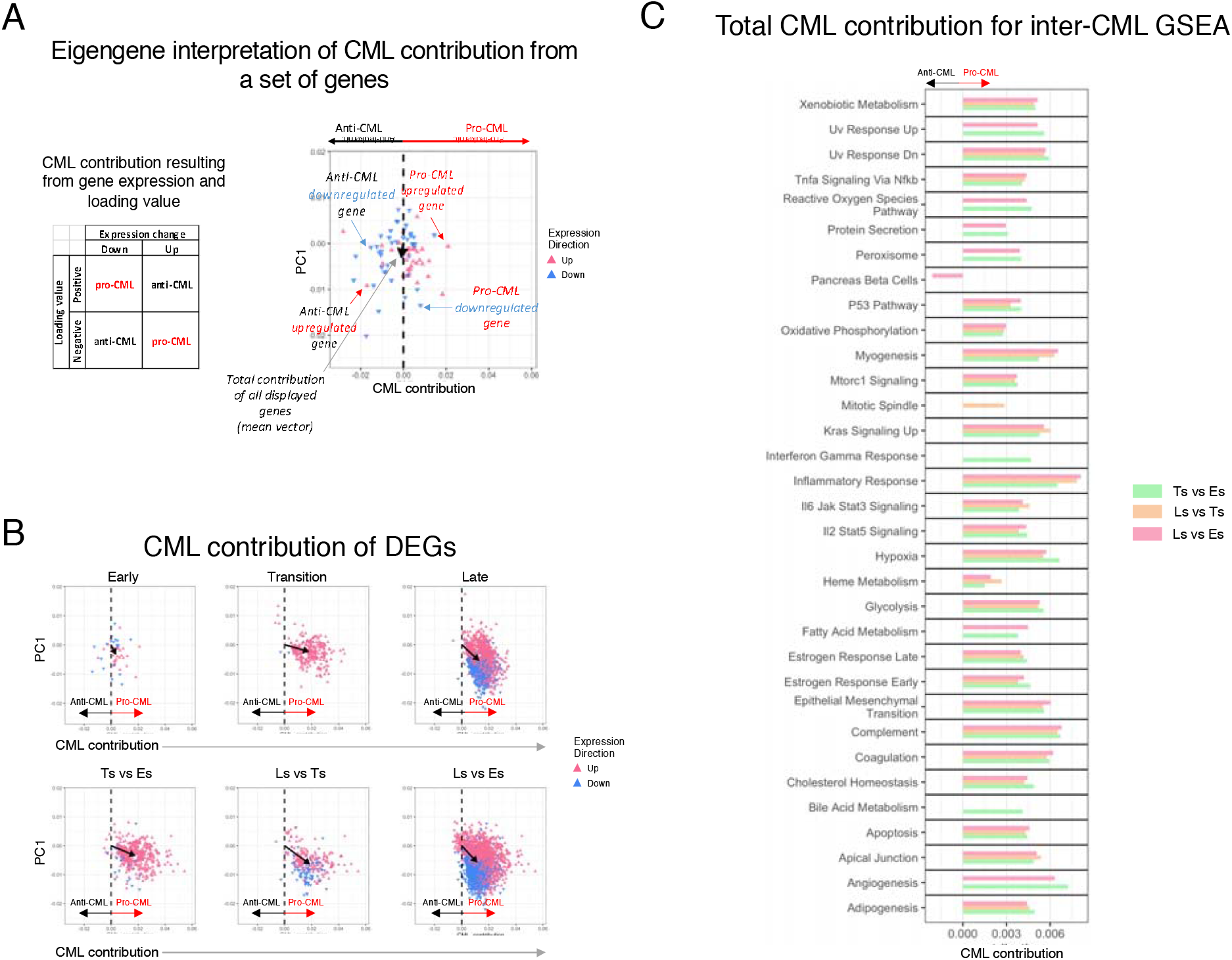
Eigengenes quantify CML contribution of genes and groups of genes. **(A)**Using the eigengenes, which were defined as the CML state-space (PC2) loading values, we determined a gene’s quantitative contribution to CML state-transition. For each gene, the CML contribution was determined by combining the eigengene with the observed gene expression change using the table (*left*) to determine whether the contribution was pro-or anti-CML. The CML contribution was illustrated for a set of genes (*right*) by plotting the CML contribution vs the PC1 value for all genes in the set. The total contribution of the set was represented by the mean vector of all genes in the set (*black arrow*). **B)** The CML contribution was used to summarize the overall effect of each DEG comparison on CML state-transition. **C)** The CML contribution was summarized for the three inter-CML comparison’s GSEA results by plotting the total CML contribution for the genes in leading edge of each significantly enriched gene set (adjusted p-value < 0.0001). Missing bars represent gene sets that were not significant in that comparison.

**Figure S5:**
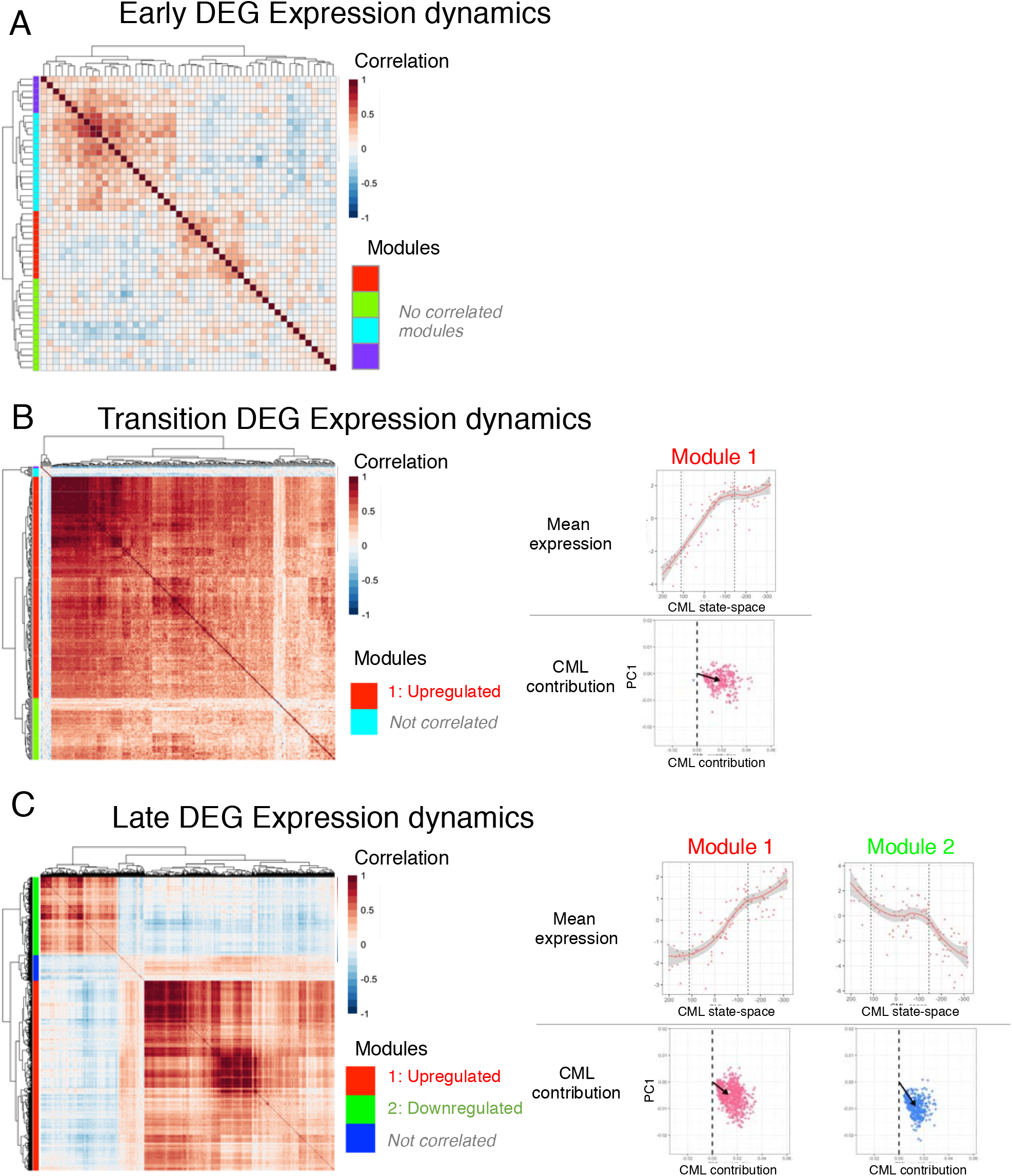
Expression dynamics of the unique disease-state DEGs. A correlation analysis using all mice from the CML cohort was used to determine which genes have similar expression dynamics over CML development. Using hierarchical clustering on the correlation matrix, gene modules were defined as a cluster of genes with a mean correlation coefficient greater than 0.25. To visualize how the expression changed over CML, the average expression of each gene module was plotted for each sample as a function of the CML statespace of the sample and summarized using a loess fit (*red line*). Finally, the overall CML contribution (*black arrow*) of each gene module was determined. This process was performed on the **A)** DEGs unique to Es, **B)** the DEGs unique to Ts, and **C)** the DEGs unique to Ls.

**Figure S6:**
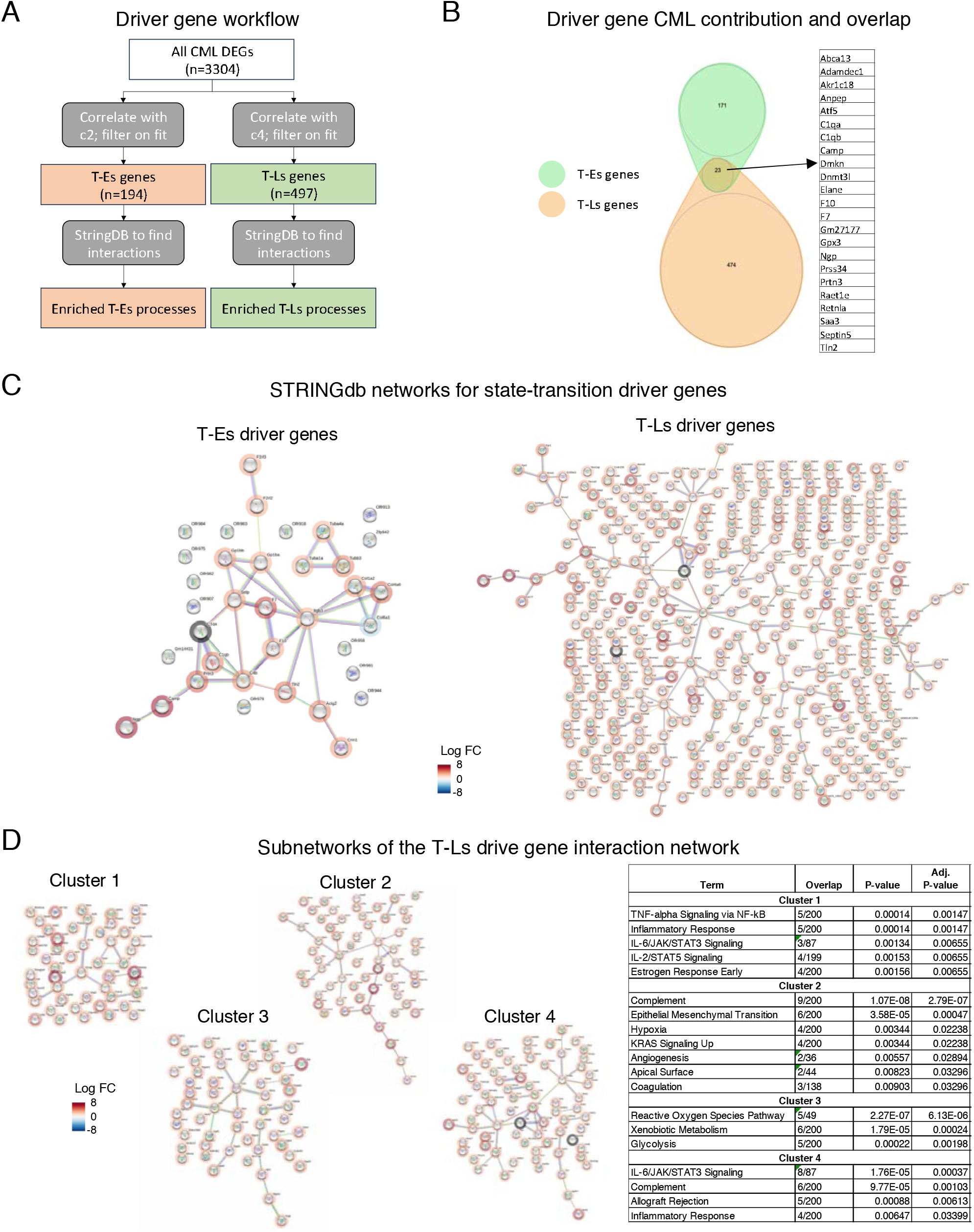
Driver gene of CML state-transition. **A)** Process for identifying the T-Es and the T-Ls drive genes. **B)** Venn diagram of the T-Es and T-Ls driver genes. **C)** Protein-protein interaction networks for the T-Es (*left*) and the T-Ls (*right*) driver genes. STRINGdb was used to identify high confidence interactions (interaction score > 900). Each gene is colored based on the observed log foldchange between Ts vs Hs for the T- Es driver genes or Ls vs Hs for the T-Ls driver genes. **D)** To further refine the processes involved in the large network of the T-Ls driver genes, four subnetworks were identified, and the significantly enriched Hallmark gene sets for each cluster were reported for each cluster (adjusted p-value < 0.01).

**Figure S7:**
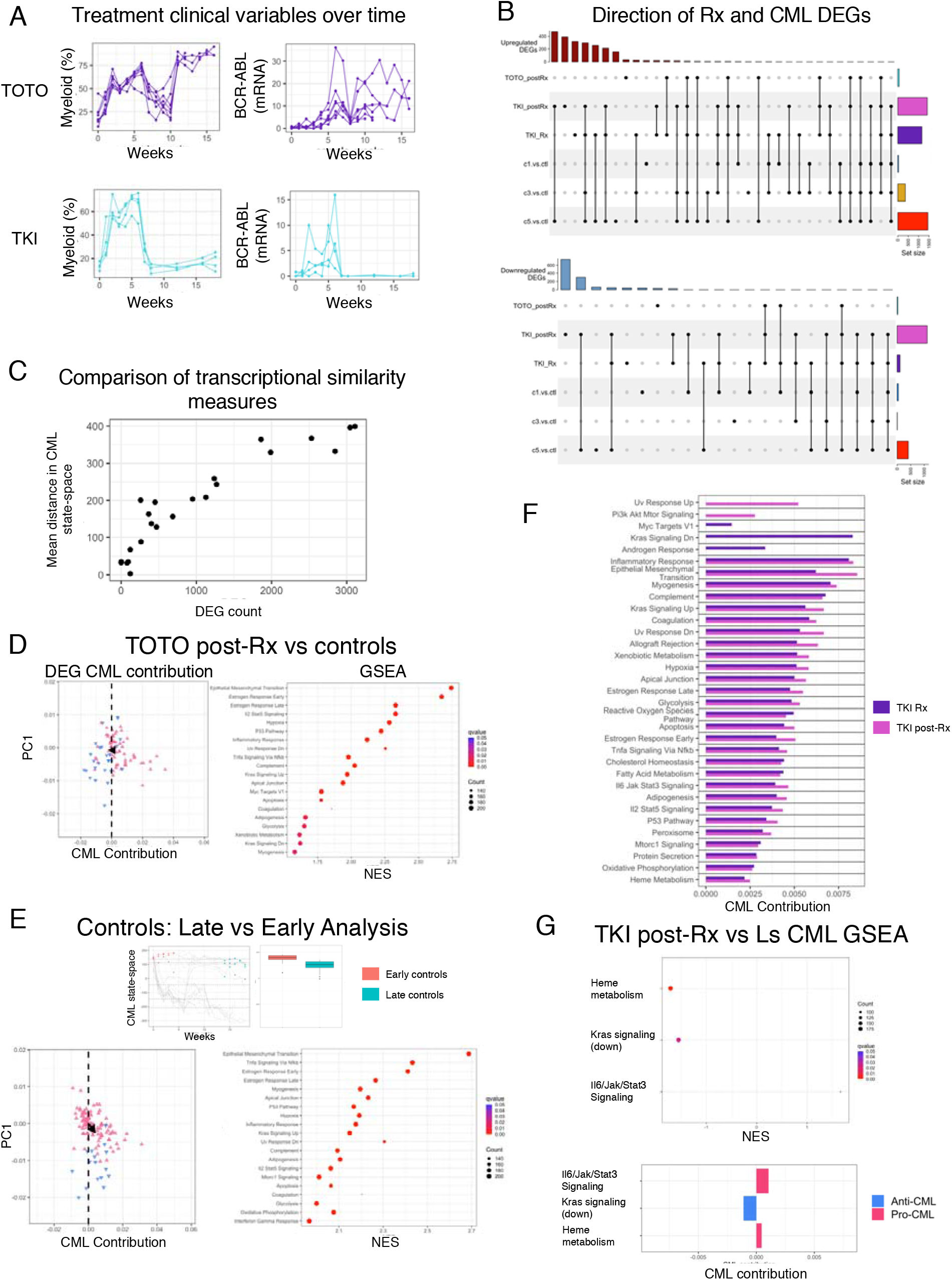
Analysis of treatment cohorts. **A)** Clinical variables plotted vs time for Tet-on Tet-off (TOTO) mice (*top*) and TKI mice (*bottom*). **B)** Intersection up- and down-regulated treatment cohorts with the disease states (Es, Ts, and Ls) compared with the control (Hs). **C)** To compare two measures of transcriptomic similarity between two groups of samples, the number of DEGs was compared to the average distance between the samples of each group in the CML state-space for all DEG comparisons. **D)** The CML contribution of the DEG (*left*) and the significantly enriched Hallmark gene sets from GSEA (adjusted p-value < 0.001; *right*) that resulted from the TOTO post-Rx samples vs healthy control (Hs) samples. **E)** An analysis comparing the early vs late control CML samples due to the observed change in CML state-space location over the course of the experiment (*top*). The total CML contribution was calculated for all DEGs that resulted from comparing the early vs late control samples (*bottom left*) which showed a small pro-CML effect in the late control mice. GSEA was also performed, and all significantly enriched Hallmark gene set pathways were upregulated in the late control mice (adjusted p-value < 0.001; *bottom right*). **F)** Total CML contribution for all significantly enriched Hallmark gene sets after GSEA for the TKI Rx and TKI post-Rx samples when compared to controls (adjusted p-value < 0.001). Missing bars indicate a gene set that was not significant in that comparison. **G)** GSEA results from the TKI post-Rx samples compared to the Ls CML samples. Significantly enriched Hallmark gene sets were shown with enrichment in expression with respect to the TKI post-Rx samples (*top*). The total CML contribution of the TKI post-Rx samples was summarized for each gene set (*bottom*).

**Table S1**: Correlation analysis of BCR-ABL expression and myeloid population with each principal component.

**Table S2**: DEG and GSEA results for both CML disease state comparisons (Es, Ts, Ls) vs healthy controls (Hs) and inter-CML disease state comparisons.

**Table S3**: Hallmark gene set enrichment analysis for state-transition driver genes at T-Es and T- Ls.

**Table S4**: DEG and GSEA results for treatment groups (TOTO post-Rx, TKI Rx, TKI post-Rx) compared to both CML disease states (Es, Ts, Ls) and healthy controls (Hs)

